# A critical evaluation of Gene Ontology priors in biologically-informed neural networks

**DOI:** 10.1101/2025.11.06.686983

**Authors:** T. Verlaan, M.A. Lieftinck, W. Mwine, M.J.T. Reinders

## Abstract

Biologically-informed neural networks (BINNs) embed prior knowledge such as the Gene Ontology (GO) into their architecture to produce interpretable representations, yet whether and how this prior improves performance or interpretation remains unclear. We introduce GONNECT, a BINN incorporating GO into an autoencoder. We evaluate GO constraints in the encoder, decoder, or both on RNA-seq tumour samples from The Cancer Genome Atlas (TCGA), comparing against published BINNs (OntoVAE, VEGA), an unconstrained baseline, randomized priors, and untrained networks. Across various performance metrics, GO structure adds little to reconstruction or latent-space organization, frequently matched by randomized or unconstrained models. Node activations agree with a gene set enrichment analysis (GSEA)-derived reference, but this agreement largely arises from the architectural constraint: it is lost when GO connections are randomized, present even before training in the encoder, and learned in the decoder. Regularized soft links outside GO (GONNECT-SL) are unstable across runs and concentrate where the ontology is sparse, compensating for the prior’s constraints rather than revealing new biology. Within this setting, we find no evidence that node activations capture biological information beyond that encoded by the predefined gene sets, suggesting that their apparent interpretability largely reflects the prior imposed by the architecture.

## Introduction

While neural networks (NNs) have proven remarkably versatile across domains^1^, their inner workings are also infamously difficult to interpret, earning them the label of “black box” models^2^. The inherent trade-off between accuracy and interpretability plays an important role in choosing the right model for a specific task^3,4^. In biological predictive modelling, understanding the mechanisms underlying functional outcomes is as important as being able to model them accurately^5,6^. Gaining insight into what the key components in a biological system are and how they interact to cause a specific phenotype is of great importance in advancing our understanding of biology and mechanisms of disease.

Earlier studies have attempted to improve the interpretability of deep learning models for biology by leveraging prior knowledge of biological systems using biologically-informed neural networks (BINNs)^7–10^, constraining the neural network structure to a hierarchical knowledge base such as the Gene Ontology (GO)^11^. Such BINNs couple each neural network node to an ontology term, and links between nodes exist only if the associated ontology terms are also linked. Due to the hierarchical nature of these ontologies, they align relatively well with the layered structure of a multilayer perceptron (MLP).

BINNs are designed to create interpretable representations of the input features, in which node activations are intended to reflect the contribution of the associated biological term. Incorporation of BINNs in prediction models allows for interpretation of these activations in the context of a predicted label, such as disease outcome or drug response^7–10^. When trained in an autoencoder setting instead, BINNs yield task-agnostic representations that are interpretable independently of any specific prediction target, and can subsequently be applied to diverse downstream tasks including classification, drug response prediction, and *in silico* modelling of gene perturbation effects^12–14^. Such autoencoders differ in the prior they impose and how it is embedded. VEGA^15^ uses a flat prior, in which a masked linear decoder ties each latent dimension to a predefined gene set, such as the Hallmark collection or Reactome, yielding a single interpretable activation per gene set but no hierarchy above it. OntoVAE^14^ instead incorporates the full GO hierarchy into its decoder, such that each decoder node corresponds to a GO term and gene expression is reconstructed through the ontology. To avoid forcing all children of a term through a single scalar, it represents each term using three neurons, which relaxes the constraint but multiplies the number of weights per edge by nine and yields three activations per term rather than one. In both models, the encoder is a conventional fully connected network.

Building a BINN on experimentally verified prior knowledge grounds the model in established biology, but that prior is incomplete. The fields it draws from remain active, so the annotations treated as ground truth are subject to extension and revision, and constraining the architecture to them can exclude relationships that have not yet been recorded. Annotation is also uneven, denser for well-studied processes and sparser elsewhere. Moreover, recent work has questioned whether it is the biological prior itself, or merely the sparsity induced by it, that drives the performance of BINNs^16^. We also observe that all existing BINN autoencoder models use prior knowledge solely in the decoder module, raising the question of whether this is preferable to a biologically-informed encoder. Although BINNs are presented as interpretable because each node maps to a biological term, it is rarely tested whether the node activations reflect the biology of that term or merely inherit it by construction. Finally, previous work addresses the incompleteness of the prior by partially relaxing its constraints, allowing the model to learn additional connections at the cost of a regularization penalty^12^. This can help a BINN overcome the rigidity and incompleteness of the prior, but raises the question of what biological meaning, if any, can be attributed to these additional connections.

In this work, we introduce GONNECT: a Gene Ontology-derived Neural Network for Explainable and Constrained Transcriptomics, and its extension GONNECT-SL, which is allowed to learn edges not present in GO at the cost of a regularization penalty. GONNECT is a BINN whose architecture is flexible in how much prior knowledge it incorporates and where, which lets us examine the questions raised earlier. We train GONNECT on bulk RNA-seq tumour samples from The Cancer Genome Atlas (TCGA), as both an encoder and a decoder, and compare it against an unconstrained fully connected MLP, two published biologically-informed baselines (OntoVAE^14^ and VEGA^15^), and randomized variants of GONNECT (GONNECT-DPR and GONNECT-FR) that retain its connectivity to varying degrees, while discarding the biological identity of its edges.

First, we find that the biological prior adds little quantitative value. Its reconstruction and clustering performance are largely matched by the unconstrained and randomized baselines, suggesting that performance depends more strongly on the connectivity structure of the prior than on the biological identity of the individual edges. Second, we find that the contribution of the prior depends on where it is placed. The GONNECT encoder governs the organization of the latent space while the GONNECT decoder governs reconstruction. Third, we assess the interpretability of the activations by comparing them to a reference derived from gene set enrichment analysis (GSEA), and find that the agreement is strongly shaped by the constraints themselves: in the encoder it is already present in untrained, randomly initialized models, whereas in the decoder it emerges after training because each node can only pass information to the genes of its term. Finally, we find that the soft links learned by GONNECT-SL improve reconstruction beyond the unconstrained baseline, but remain unstable across initializations and concentrate in the sparsest regions of the ontology, so they reflect a response to ontology sparsity rather than evidence for specific biological relationships. Taken together, the interpretability of these models largely reflects the prior they are given rather than biology they uncover.

## Results

### GONNECT

We present GONNECT, a Gene Ontology-derived Neural Network for Explainable and Constrained Transcriptomics. We use the hierarchical prior knowledge in Gene Ontology (GO) as GONNECT’s architecture (Figure 1a). Before use in models, the GO ontology graph is filtered and processed into a graph that is suitable as a neural network architecture (Methods); this processing determines the number of layers, nodes and weights of GONNECT (Figure 1b). With our settings the processed graph comprises 623 GO terms, 2,369 proxy terms, and 1,000 genes. Consequently, the resulting GONNECT architecture has an input dimension of 1,000, five layers, and an embedding dimension of 109.

**Figure 1.**
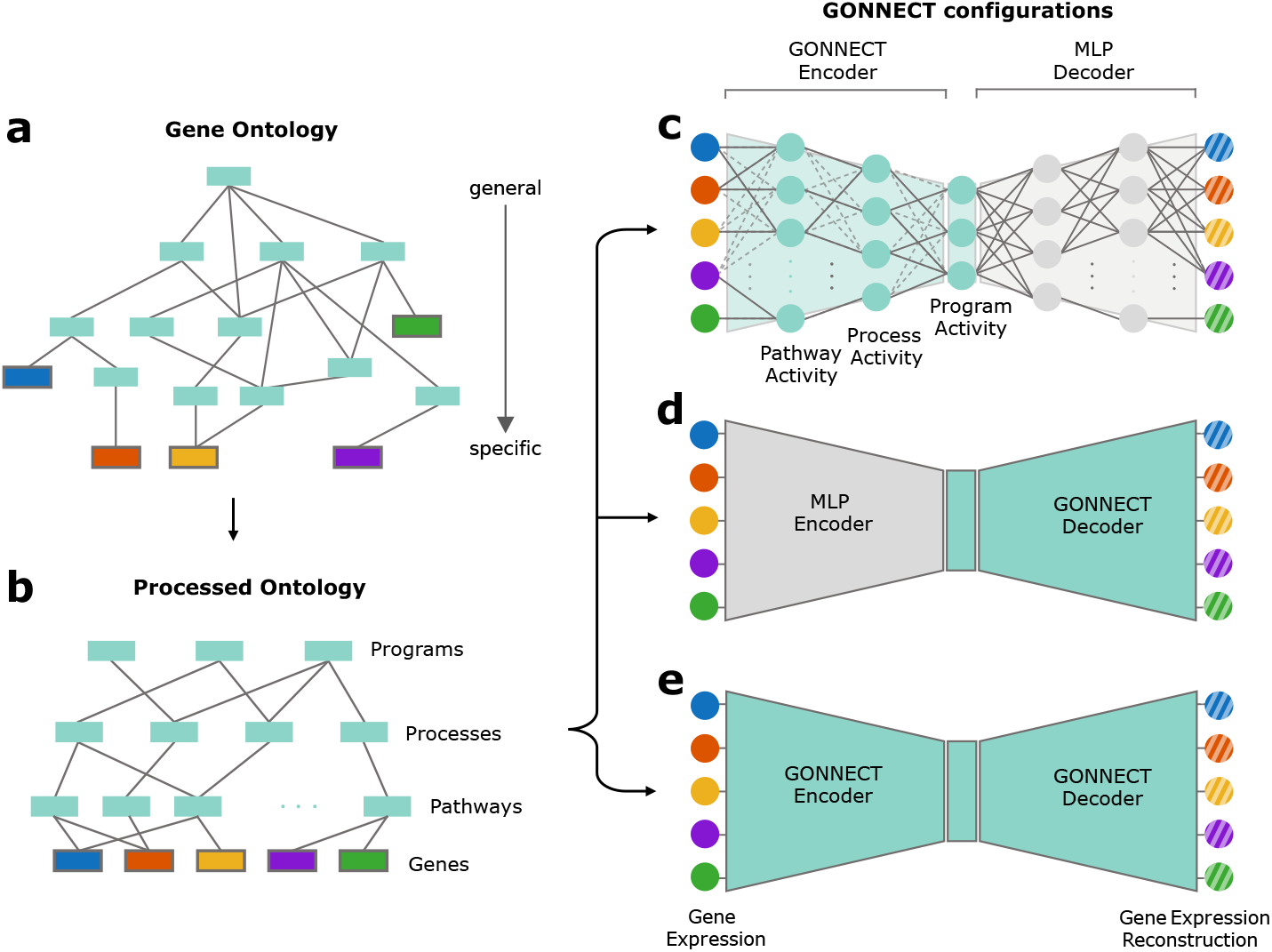
Overview of GONNECT. **a**) A hierarchical ontology graph containing experimentally verified prior knowledge on biological entities (GO terms) and their relationships. Boxes are GO terms; the coloured nodes attached to them are the genes annotated to those terms. **b**) Processed GO that can be used as a sparse neural network architecture. The required structure is achieved by pruning and condensing parts of the network that are too sparse, and then balancing it using proxy terms (Methods for details). Note that GO does not actually include the “programs”, “processes” and “pathways” annotations.They are just there to illustrate the hierarchical nature of the ontology. **c**) Autoencoder with a GONNECT encoder such that network nodes represent GO terms and solid links represent known relationships between terms. The decoder is a fully connected MLP with an identical but mirrored node layout. The inputs to the network are the observed gene expression levels in a sample (coloured nodes on left), which are reconstructed by the autoencoder (shaded colour nodes on the right). Dotted lines represent the soft links that can be added by GONNECT-SL to complement biological prior information from GO. **d**) Configuration where the encoder is an MLP and the decoder is a GONNECT module. **e**) Configuration where both encoder and decoder are GONNECT modules, such that every node in the network is coupled to a GO term.

Biology can be incorporated in GONNECT through three different configurations: one in which a GONNECT encoder is coupled to a biologically-agnostic, fully connected MLP decoder (Figure 1c), a second configuration consisting of an MLP encoder and GONNECT decoder, in which the GONNECT architecture is a mirrored version (Figure 1d), and a third in which both the encoder and decoder are GONNECT architectures (Figure 1e). In all three configurations, the embedding space is coupled to a GONNECT architecture, meaning that nodes in the embedding space are associated with GO terms. We also assess GONNECT-SL, a version of GONNECT where instead of fixing the weights of connections that do not exist in GO to zero, we apply a weight regularization (Methods). This allows the model to learn soft links, which increases model flexibility at the cost of a penalty.

To assess the performance of GONNECT, we use the publicly available The Cancer Genome Atlas (TCGA)^17^ dataset, which contains gene expression from human tumour samples from many different cancer types. After preprocessing, the dataset contained 9,797 samples, from 32 different cancer types, and their 1,000 most highly variable genes with at least one GO-annotation (Methods). The distribution of cancer types in the preprocessed dataset is available in Supplementary Table S1.

### Encoder and decoder respectively dictate latent structure quality and reconstruction performance

We compared GONNECT to a biologically-agnostic reference, a fully connected MLP with the same number of layers and nodes per layer as GONNECT. Since each model constrains one module and leaves the other fully connected, we compare module against module: a dense module contains 3,360,429 learnable weights, compared with 3,138 for a GO-structured encoder and 4,100 for a GO-structured decoder. We also compared GONNECT with the two previously published biologically-informed autoencoders described in the introduction, OntoVAE and VEGA, both of which place their prior in the decoder. OntoVAE applies its own, less extensive processing to GO, retaining 4,023 terms against our 623. With three neurons per term, its decoder is approximately one hundred times larger than GONNECT’s. VEGA’s masked decoder is smaller than GONNECT’s with 867 learnable connections when using the Hallmark gene sets as prior and 1,984 when using Reactome pathways (see Methods for more details on OntoVAE and VEGA). All models and variants are evaluated over the same five independent training runs, each with its own random data split (Methods).

When we consider the average reconstruction loss (Figure 2a, mean squared error), the model with biology only in the encoder performs on par with the MLP and better than both VEGA variants, but is outperformed by OntoVAE. Remarkably, the GONNECT decoder model, which like VEGA and OntoVAE incorporates biological information into the decoder, performs significantly worse. This asymmetry arises from the hierarchical structure of GO: an *encoder* node aggregates information from multiple children into a single activation while preserving flexibility, whereas a *decoder* node requires multiple children to share a single parent activation, thereby leaving their representation entangled. Neither baseline pays this cost in the same way. OntoVAE widens the parent bottleneck by using three neurons per term, which, combined with the larger ontology it retains, increases the number of learnable weights by approximately 100-fold and yields three activations per GO term, thereby complicating interpretation. VEGA has no parent layer at all, as its flat prior spans a single masked layer from gene sets to genes.

**Figure 2.**
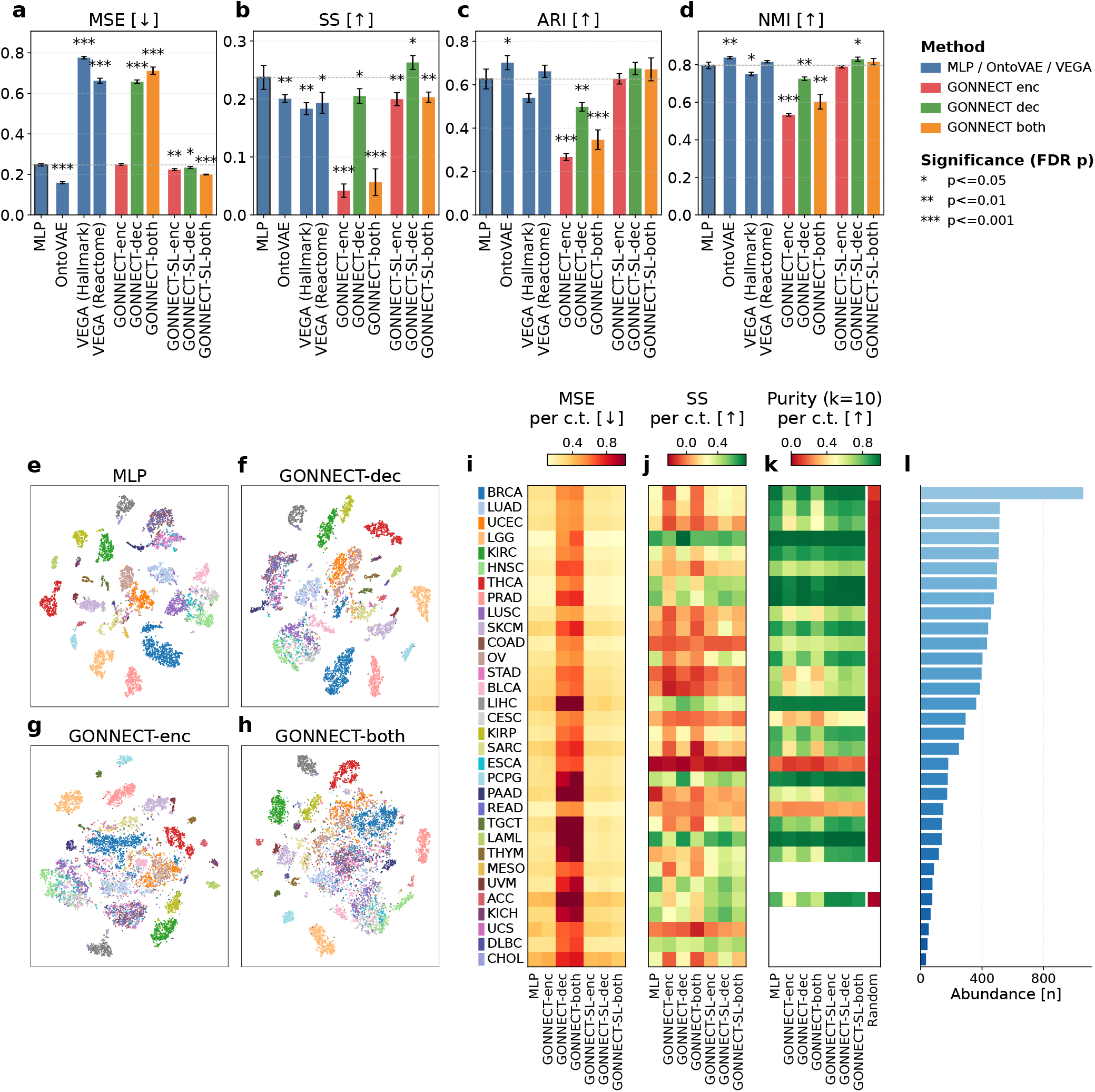
Performance metrics and t-SNE visualizations for different GONNECT variants and a comparison to baselines. **a-d**) mean squared error (MSE), silhouette score (SS), adjusted Rand index (ARI) and normalized mutual information (NMI) for all model variants, computed on the held-out test samples. Colours indicate where the GONNECT modules sit (‘enc’ for encoder, ‘dec’ for decoder, ‘both’ for both). Bar heights denote the mean over five independently trained model instances and error bars the standard deviation; arrows indicate whether higher or lower values are better. The dashed line indicates the MLP reference; asterisks indicate a significant difference from it in either direction. **e-h**) t-SNE visualizations of latent spaces learned by the MLP reference model, GONNECT-dec, GONNECT-enc and GONNECT-both, coloured by cancer type (soft-link equivalents in Supplementary Figure S2). **i-k**) MSE, SS and *k*-nearest-neighbour purity per cancer type (‘c.t.’) for each main model, sorted by cancer type abundance (panel l). Purity is the fraction of a test sample’s *k* = 10 nearest test-set neighbours carrying its own cancer type, averaged per type and over the five instances; types with fewer than *k* test samples in any of the five splits are not shown, and the rightmost column shows the purity expected for randomly drawn neighbours (Methods). Supplementary Figure S3 shows additional *k* settings and models.

Next we consider several clustering metrics. The silhouette score (SS) assesses geometry by comparing intra- to inter-cluster distances, whereas ARI and NMI measure agreement between a k-means clustering and the true cancer-type labels (Figure 2b-d). This shows that the gap between the reference models and the GONNECT decoder largely closes, indicating that latent organization is less sensitive to the sparsity of a biologically-informed decoder. The GONNECT encoder, however, now shows a large drop, also visible in the t-SNE visualizations (Figure 2e-h), where the MLP and GONNECT decoder result in neatly organized, well-separated cancer type clusters, while the GONNECT encoder does not. Reconstruction and latent organization are therefore determined by different modules: i.e. reconstruction is governed by the decoder, whereas latent organization is governed by the encoder. This also explains GONNECT-both, which tracks the weaker of the two individual BINNs on every metric (Figure 2a-d). The effect of the biological constraint therefore remains local to the module in which it is placed. The soft-link variants (GONNECT-SL) bring all three configurations to a lower MSE than the MLP and bring ARI and NMI close to the MLP reference; only the silhouette score of GONNECT-SL-enc and GONNECT-SL-both remains below the reference (Figure 2a-d). Their latent spaces are shown in Supplementary Figure S2.

We next examine reconstruction and embedding quality per cancer type (Figure 2i-k). Reconstruction difficulty was largely a property of the cancer type for the MLP, the GONNECT encoder and the soft-link models, which reconstructed the same types poorly, whereas the constrained decoders struggled most with a different set of types (e.g. LAML, TGCT and THYM) on top of the overall decoder effect observed in panels a-d. This per-type difficulty partially tracks sample abundance (Figure 2l), with rare cancer types such as ACC, UCS and CHOL showing among the highest reconstruction errors. The silhouette score shows no such trend and is lowest for digestive-system types (ESCA, PAAD, COAD and STAD). Cancer types from the same tissue of origin overlap in the t-SNE (Figure 2e-h), with digestive-system types (ESCA, STAD, COAD and READ) and female reproductive-system types (UCEC, UCS, CESC and OV) each falling in a shared neighbourhood. These tissues share large parts of their expression profiles, so the embedding places them close together and the clustering metrics treat the residual within-tissue differences as harder to resolve, lowering the neighbourhood purity of ESCA and READ in particular (Figure 2k). Every cancer type shown nevertheless retains a purity well above that expected for randomly drawn neighbours (Figure 2k, rightmost column). Supplementary Figure S3 repeats these per-type metrics for all ten model variants and for *k* = 10, 20 and 30.

The processing steps described above include several free parameters that shape the resulting topology. We therefore varied the child-term threshold, the minimum layer width and the skip-connection pruning, with the parent threshold fixed at 1, recording the topology for every combination (Supplementary Table S2). With skip-connection pruning, the child-term threshold had the largest effect on the network topology, with higher values producing a coarser ontology containing fewer, more aggregated terms; the minimum layer width changed little between 20 and 50 but strongly reduced depth at 200. Without skip-connection pruning, about five times as many terms were retained. We therefore varied the child-term threshold (ct=5, 10, 30) at 1,000 input genes, and separately compared 1,000 against 2,000 input genes at ct=30, while holding the remaining parameters at their defaults (Supplementary Figure S1). Doubling the number of input genes roughly doubled the number of retained GO terms, reflecting that ontology pruning depends on the set of genes retained during feature selection. Performance was largely stable across all these settings, and the configuration used in the main text (ct=30, 1,000 genes) performed at or near the best of those tested for most models. There were two exceptions. The GONNECT encoder separated the cancer types better at ct=5 and ct=10. The GONNECT decoder (GONNECT-dec), in contrast, reached a reconstruction error at ct=5 and ct=10 approximately equal to that of predicting each gene’s mean (MSE ≈ 0.99), and its silhouette score collapsed, while its ARI and NMI changed little. A finer-grained decoder therefore yields an embedding whose clusters are still correctly assigned, as reflected by ARI and NMI, but are no longer compact or well separated in distance, as reflected by the lower SS.

In terms of training time, the models require more or less the same time per epoch, but differ in the number of epochs until convergence (Supplementary Table S3). The GONNECT encoder converged in about as many epochs as the MLP, whereas GONNECT-dec required about 4.5 times as many, and GONNECT-both at least 20 times as many. We attribute this to the constrained connectivity of the GONNECT decoder, in which substantially fewer paths connect successive layers for gradient propagation than in the fully connected MLP, and through which the gradients for the whole network must pass before reaching the encoder. With soft links, GONNECT-SL-enc and GONNECT-SL-dec converged in about 1.3 and 1.1 times as many epochs as the MLP, and GONNECT-SL-both in about 3.6 times as many.

### BINNs and randomized equivalents perform similarly

To test whether GONNECT’s performance derives from the specific biological identity of its connections or merely from their connectivity structure, we compared every model against two randomized variants of its own graph: a degree-preserving variant GONNECT-DPR, in which edges are rewired while holding each node’s in- and out-degree fixed (Methods), and a fully randomized variant, GONNECT-FR, where the number of nodes and edges per layer are retained, but the connectivity is fully random. The two variants therefore distinguish three possibilities: that performance derives from the biological identity of the edges, from the degree structure and localized sparsity imposed by the processed ontology, or from sparsity alone. GONNECT-DPR removes the first while retaining the second, and GONNECT-FR removes the second as well.

Across models and metrics, performance changed little under either randomization. For most models, GONNECT-DPR was not significantly different from the original graph on silhouette score, ARI and NMI (Figure 3b, Supplementary Figure S4). On MSE, degree-preserving randomization increased reconstruction error in five of nine models, by 6–13% for the three GONNECT configurations, and decreased it for OntoVAE (Figure 3a). However, the two randomized models differ systematically. GONNECT-DPR rarely improved upon the unrandomized GONNECT, whereas GONNECT-FR, where it had an effect, tended to improve reconstruction beyond GONNECT-DPR and, in some cases, the unrandomized model itself. We interpret this as a consequence of what each randomization method removes. Degree-preserving randomization discards the biological identity of the edges but retains the localized sparsity and degree structure of the processed ontology, thereby preserving the information bottleneck and, consequently, allowing performance to at best match that of the original graph. Full randomization additionally relaxes the degree structure and thus the bottleneck in low-degree regions of the network, permitting more flexible information flow. These results suggest that, under the conditions examined here, performance depends more strongly on the connectivity structure of the processed ontology than on the biological identity of individual edges.

**Figure 3.**
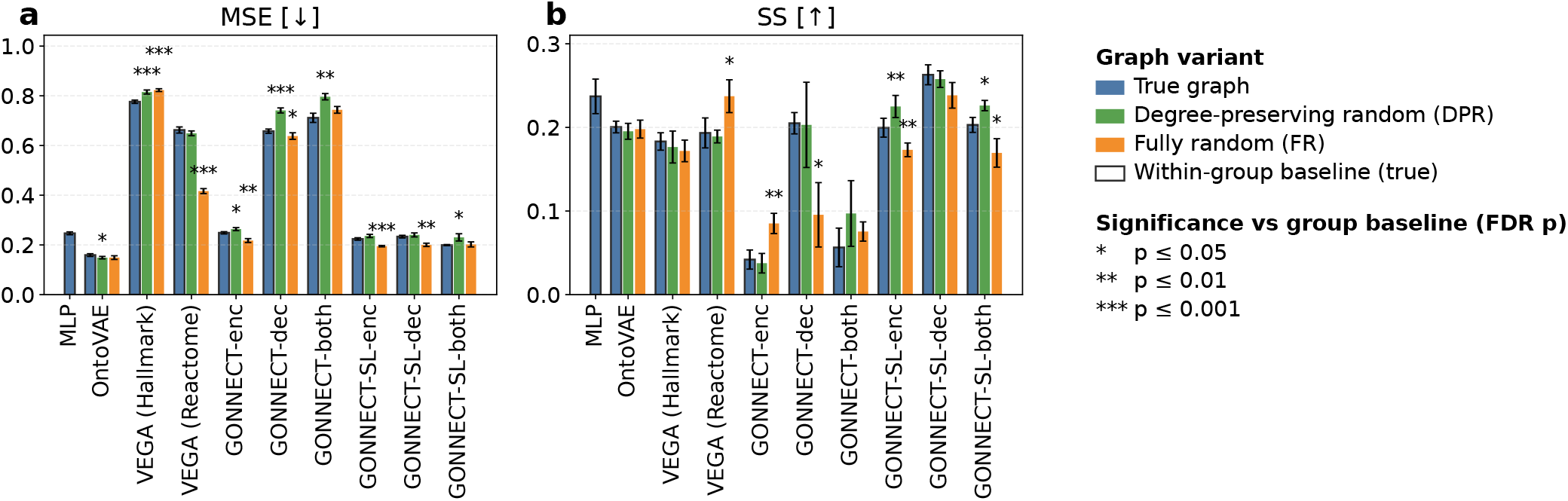
Comparison of each model with two randomized alternatives: (i) a degree-preserving model where each term keeps its degree, but connectivity to other terms is randomized, and (ii) a fully randomized model where only the number of edges per layer is preserved. Significance is reported with respect to a model’s unrandomized equivalent. We report mean squared error (MSE, panel **a**), and silhouette score (SS, panel **b**), both computed on the held-out test samples. See Supplementary Figure S4 for ARI and NMI, which show patterns highly similar to SS.

A few informative cases deviated from this general pattern. The clearest example concerns the silhouette score of the GONNECT encoder and decoder, which responded in opposite ways to full randomization. The encoder, which had the lowest silhouette score of all models, improved under full randomization, whereas the decoder, among the highest, deteriorated. In contrast, the degree-preserving randomization left both modules close to their performance on the original graph. One possible explanation is that the embedding is shaped primarily by the degree-constrained bottleneck imposed by the processed topology, rather than the biological identity of the individual edges, with opposite effects on the two modules. The GONNECT encoder achieves lower SS than the MLP, suggesting this structural constraint hinders its embedding. Full randomization may therefore relax the constraint and improve the metric. The decoder, by contrast, already produces a strong embedding despite this structure, so full randomization may instead impair performance by disrupting the degree organization that supports it. Consistent with this interpretation, degree-preserving randomization closely matches the original graph for both modules, indicating that the observed effects arise from the overall connectivity structure rather than from the specific biological relationships encoded by the edges.

The soft-link models exhibited a more consistent pattern: degree-preserving randomization generally increased reconstruction error, whereas full randomization tended to reduce it, with the embedding metrics changing in the opposite direction. These effects were small and not consistently significant. They were most evident in the silhouette score; ARI was unchanged and NMI shifted only slightly, in the same direction (Supplementary Figure S4). We therefore interpret them as modest changes in cluster compactness rather than a broad reorganization of the embedding.

### Node activations agree with gene-set enrichment by construction

Having established that biological information is not necessarily beneficial in terms of reconstruction or latent clustering performance, we now examine whether the node activations of the biologically-informed models make sense from a biological point of view. We consider the mean activation of each GO term per cancer type and find that over repeated training experiments, node activation magnitudes per cancer type stay similar in the encoder (Supplementary Figure S5), but much less so in the decoder (Supplementary Figure S6). This indicates that the GONNECT encoder repeatedly treated the same GO terms as active for a given cancer type. The sign of these activations, centred per GO term in Supplementary Figures S5 and S6, may differ between model instances because of random weight initialization and the unconstrained weight signs during training. Importantly, these signs cannot be interpreted as indicating positive or negative biological regulation, because the GO *is_a* relations used to construct the network encode subtype relationships between terms rather than regulatory direction. We therefore summarize each node by the ROC-AUC of a classifier predicting cancer type from the GO term activations, yielding an unsigned association score for each “cancer type–GO-term” pair (Supplementary Tables S4 and S5).

To assess whether the term activations reflect the biology of their terms, we compared them with a reference derived from gene set enrichment analysis (GSEA). For each cancer type, we identified genes differentially expressed relative to all other cancer types and quantified their enrichment within the set of genes annotated to each GO term or its descendants in the processed GO graph (Methods). In the GONNECT encoder, term activations in each layer were significantly correlated with the GSEA-derived reference, whereas in the decoder the bottleneck layer (L5) and, only weakly, L6 and L7 reached significance (Figure 4a). GONNECT-SL showed significant but lower agreement in encoder layers L0–L2, and none at the bottleneck (L3) or in any decoder layer (Figure 4b). For the OntoVAE, we found significant correlations in 8 of the 11 layers; only L1, L9 and the final layer (L10) were not significantly correlated (Figure 4d). VEGA’s activations were significantly correlated for both prior knowledge bases, Hallmark and Reactome, but the correlations were stronger for Hallmark, which has more edges per gene set than Reactome (Figure 4e). For the randomized models from the previous section, we observed no significant correlations (Figure 4c,e), as expected since the gene–term relations on which the comparison depends were broken during randomization. Overall, the GONNECT encoder layers agreed more strongly with the reference than the decoder layers.

**Figure 4.**
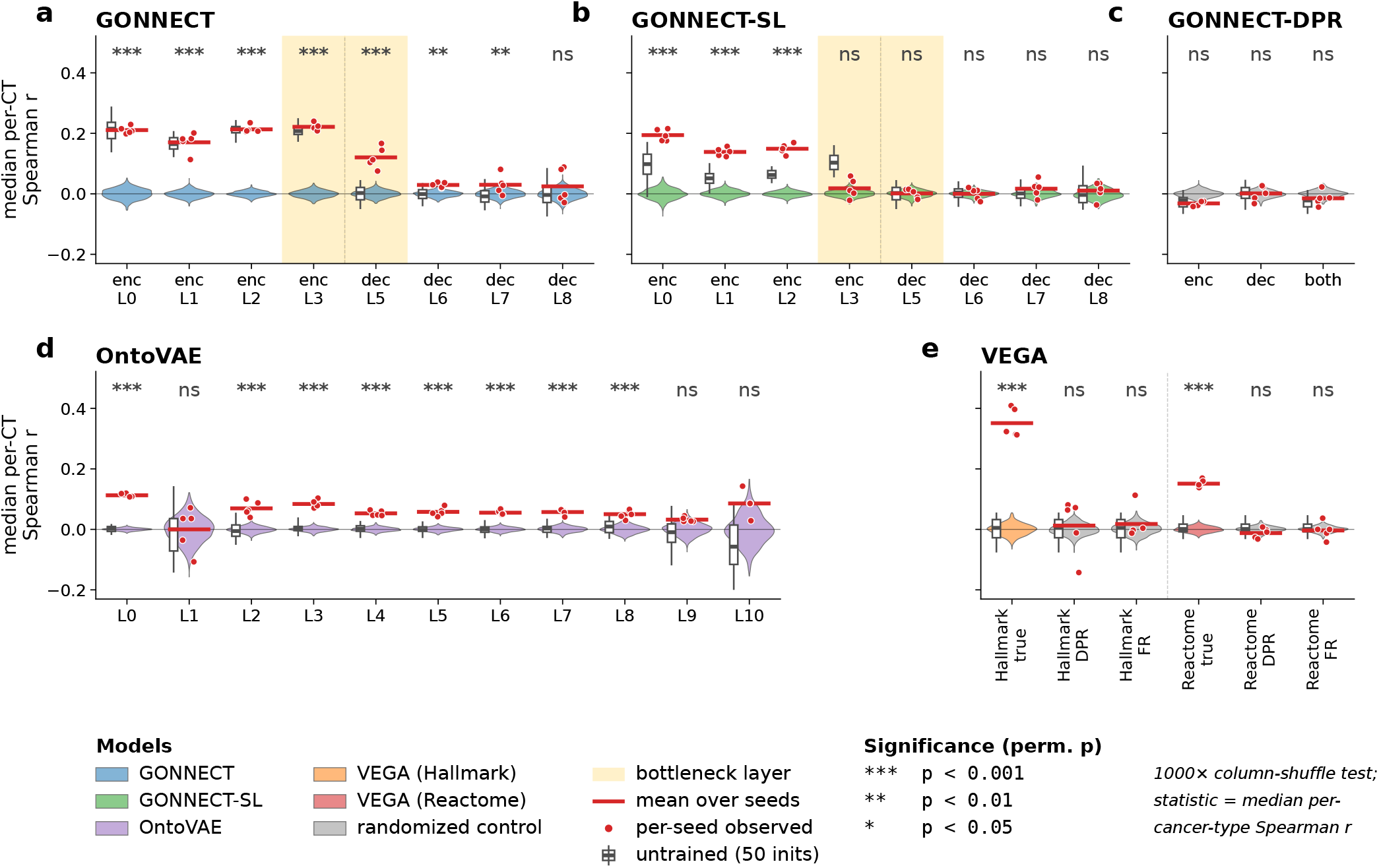
Activation–enrichment agreement across models and layers. Per layer we show the median per-cancer-type Spearman correlation between node cancer-type specificity (measured by ROC-AUC) and gene-set enrichment (− log_10_ GSEA *p*-value, see Methods), computed separately for each seed (red dots); the red line is the mean over five seeds. The violins show the distribution of this mean under a permutation test, where GO terms are randomly shuffled, and the stars give its one-sided significance (^∗^ *p <* 0.05, ^∗∗^ *p <* 0.01, ^∗∗∗^ *p <* 0.001). The small boxplots show the same statistic for untrained, randomly initialized models (50 random initializations per model, each scored separately), giving the agreement present before any training. Where the untrained activations are computed by the fully connected encoder alone (decoder L5 in **a** and **b**, and VEGA in **e**), they do not depend on the ontology wiring, and the boxes coincide. In **c**, the untrained encoder and both models are initialized with the same seeds and therefore share an identical encoder, so their boxes coincide as well. **a)** original GONNECT and **b)** GONNECT-SL, with the encoder layers (L0–L3) taken from the encoder models (GONNECT-enc, GONNECT-SL-enc) and the decoder layers (L5–L8) from the decoder models (GONNECT-dec, GONNECT-SL-dec), and the 109-node latent (encoder L3 / decoder L5) highlighted; **c)** GONNECT-DPR, with the degree-preserving randomized ontology; **d)** OntoVAE layers L0–L10; **e)** VEGA, which has only one decoder layer (Hallmark, Reactome) with true, degree-preserving and random ontologies. Randomized-ontology controls are shown in grey.

Part of this agreement is expected from the architecture, because each encoder node combines the expression of the genes annotated to its term. We therefore repeated the analysis on untrained, randomly initialized models (Methods). In the encoder, untrained and trained models agreed with the reference to the same degree, with median correlations of 0.17–0.22 both before and after training across L0–L3 (Figure 4a), so this agreement follows from the architecture rather than from training. With the degree-preserving randomized ontology, agreement was close to zero both before and after training (Figure 4c), so this component requires each node to be wired to the genes of its term, rather than following from the connectivity structure alone. The permutation test, which shuffles term identity, does not capture this component. In GONNECT-SL, whose soft links start as small random connections, untrained agreement was lower, and training raised it in L0–L2 but lowered it at the bottleneck (Figure 4b). In the decoder, the architecture is likewise constrained because each node can only pass information to the genes of its term, but the untrained control cannot directly quantify the resulting agreement: a decoder node’s activation is computed by the upstream network, including a fully connected encoder that is untrained as well, so it carries no information about its term before training. Accordingly, untrained agreement was close to zero in the GONNECT decoder, OntoVAE and VEGA (Figure 4a,d,e), so the agreement in these decoder-based models, including the significant correlations in most OntoVAE layers and for both VEGA gene-set collections, arose during training. The trained decoder with the degree-preserving randomized ontology, whose nodes pass information to other genes than those of their term, showed no significant agreement (Figure 4c), so the agreement acquired during training likewise requires each node to be wired to the genes of its term.

### Soft links improve reconstruction while retaining much of the prior structure

GONNECT-SL relaxes the strict GO constraint by allowing soft links, weights between terms that are not connected in the original ontology, at the cost of a regularization penalty. These soft links closed the reconstruction gap that the original GONNECT showed in the GONNECT-dec and GONNECT-both configurations, bringing all three configurations to a lower MSE than the MLP baseline, although the silhouette score of GONNECT-SL-enc and GONNECT-SL-both remains below that of the MLP (Figure 2a–d). We selected the regularization strength α from a sweep from 10^2^ to 10^5^ over reconstruction performance and the number of active soft links (Supplementary Figures S7 and S8). Increasing α suppresses the soft links and moves the model towards the original GONNECT: in the decoder, 635 soft links are active at α = 10^2^, 86 at α = 10^3^, two at α = 10^4^ and none at α = 10^5^, where the relaxed model has collapsed onto the original ontology. Its reconstruction error rises from 0.205 to 0.649 across the same range. We chose α = 1 · 10^2^, the only value at which the soft-link decoder recovers the reconstruction of the fully connected models. As for all other configurations, these models were trained with early stopping (Methods).

The soft links complement the ontology rather than overriding it. The weights of links already present in GO retained a distribution close to that of original GONNECT (Figure 5a), indicating that GONNECT-SL did not learn to suppress known edges. The non-GO soft-link weights remained below the GO-edge weights, with their bulk at least an order of magnitude below the strongest GO edges, so they did not dominate the learned weights.

**Figure 5.**
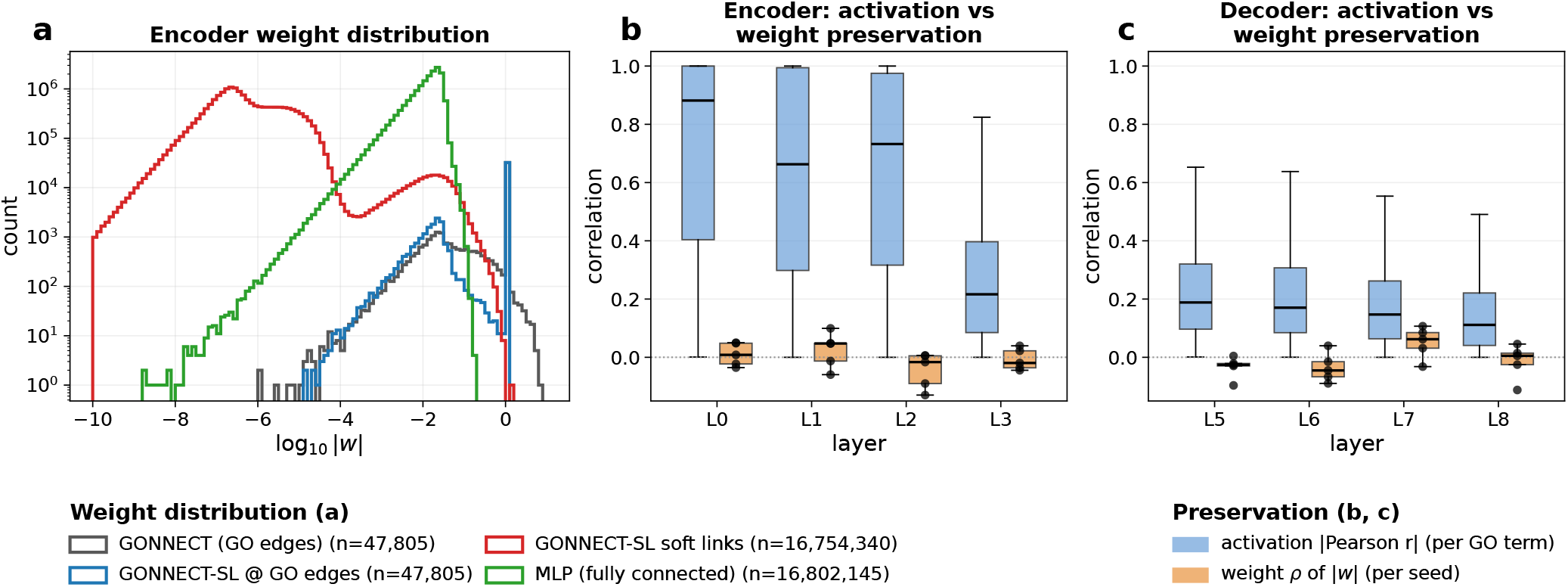
a) Distribution of learned (log absolute) weights of the GONNECT-SL encoder, pooled over the five seeds. For comparison, the weight distributions of a fully connected MLP encoder and of the GONNECT encoder are shown. The weights of GONNECT-SL are split up into the weights of links that were already present in GO and those that are not. The spike at 1.0 represents the fixed weights of the proxy terms introduced in the GO processing (Methods), which are included in the GO-edge counts. **b,c**) Activation vs weight preservation between GONNECT and GONNECT-SL, per layer, for the encoder (L0–L3, panel **b**) and the decoder (L5–L8, panel **c**). Blue: per-GO-term activation similarity (|Pearson *r*| of GONNECT vs GONNECT-SL activations over all 9,797 samples, same seed). Orange: agreement of the GO-edge weights themselves (per-seed Spearman ρ of |*w*| over biological GO edges; dots = 5 seeds).

To test whether the relaxation preserves the biological signal captured by the fixed ontology, we compared GONNECT-SL activations to those of original GONNECT. Per-node activations were most strongly preserved in the first encoder layer, closest to the genes, and declined with depth (Figure 5b). The decoder showed the same decline along the direction of information flow: correspondence was highest in the layer closest to the bottleneck and decreased monotonically towards the gene-level output layer (Figure 5c). The GO-edge weights themselves were essentially uncorrelated between the two models (Figure 5b,c).

The weight magnitudes of the soft links were moderately rank-correlated across random seeds (pairwise Spearman ρ≈ 0.53 for the encoder and ≈0.66 for the decoder; Supplementary Figure S9), but the identities of the strongest links were far less stable. Of the links we classified as “active” (|*w* |*>* 0.1), the large majority were active in only one of the five seeds, and only a small set were active in all five (32 in the encoder, none in the decoder; Supplementary Figure S9). Together this indicates that the biological prior was retained most strongly in the gene-proximal encoder layers, but that each model instance retained it using a different combination of weights.

Next, we considered whether the learned soft links may have a biological meaning to them. To assess this, we held out a random 10% of GO edges and asked whether GONNECT-SL re-learned them as high-weight soft links more often than random non-GO edges. Held-out GO edges were not recovered preferentially (Supplementary Figures S10 and S11). Recovery was mainly associated with the density of GO terms in the neighbourhood of the term, rather than with its held-out status. After controlling for GO density (measured by out-degree in the ontology) we found that in a few cases held-out GO links were recovered significantly more often, but not consistently enough to consider soft-link placement biologically meaningful.

We also considered whether soft links appeared where the ontology was most heavily modified during processing, by relating each soft-link weight to the number of merge and prune operations in the associated nodes’ subtrees and supertrees. Across most layers, we found little relationship (Supplementary Figure S12). Only in the sparsest layer of the decoder did we observe a positive correlation between soft-link weight and the number of merge and prune operations in the subtree (Spearman ρ up to ≈ 0.4). This is consistent with soft links concentrating in the sparsest parts of the graph, but overall, soft-link weights were largely independent of where the ontology was merged or pruned.

Taken together, the soft links appeared predominantly in the sparse regions of GO, where the autoencoder’s information bottleneck is tightest, and their individual identities were unstable across repeats. We therefore regard them as a means of addressing ontology sparsity rather than as a reliable mechanism for proposing novel biological relationships. At the same time, the activations produced by the relaxed model still corresponded substantially to those of the original model in the gene-proximal encoder layers, indicating that the approach retains much of the correspondence imposed by the prior where the data constrains it most.

## Discussion

We introduced GONNECT, a sparse, biologically-informed neural network that leverages prior knowledge from the Gene Ontology with the aim of making the latent space of gene expression autoencoders interpretable in terms of biological processes and their relationships. By mapping each neuron to a GO term and wiring edges to match GO relationships, GONNECT produces a network in which every node is labelled by a GO term, and because it can serve as encoder, decoder, or both, it lets us isolate what the GO scaffold contributes in each role. Rather than treating the biological prior as inherently beneficial, we used this flexibility to assess what it contributes quantitatively. Within the setting examined here, the gains over matched baselines are limited, which leaves interpretation as the main potential value of the approach. Node activations do agree with a gene-set enrichment reference, but this agreement is strongly shaped by the constraints that tie each node to the genes of its term, and we find no evidence that the activations reflect biology beyond these gene sets.

Comparison between the encoder and decoder versions of GONNECT shows that reconstruction quality is mainly determined by the decoder, and latent organization by the encoder. The encoder matches a fully connected MLP on reconstruction and exceeds both VEGA variants, though not OntoVAE. The decoder-side prior degrades reconstruction but performs more similarly to the baseline on clustering metrics. This shows that the effect of the constraint stays local to the module it is placed in. No placement improves every performance metric at once.

Through randomization of the models, we find that the biological identity of the GO edges has little bearing on performance. Replacing the true edges with degree-preserving randomized versions, which keep each node’s in- and out-degree but discard biological identity, left the clustering metrics unchanged in most cases and increased reconstruction error only modestly. The observed performance is therefore determined mainly by the connectivity structure of the processed ontology rather than the biological identity of the edges. This concurs with recent work finding that the sparsity pattern of biologically-informed networks, rather than their biological content, largely drives their performance^16^.

The biological identity of the edges contributes little quantitatively, and the node activations offer less biological insight than their labels suggest. In the encoder, the activation magnitude of each GO term per cancer type is stable across repeated training runs, indicating that the GONNECT encoder consistently treats the same terms as active for a given cancer type. To test whether these activations reflect the biology of their associated terms, we compared per-node cancer-type specificity against gene-set enrichment for the same term, and find that the two align significantly in every encoder layer and more weakly, in three of four layers, in the decoder.

This alignment is strongly shaped by the architectural constraints. An encoder node combines the gene inputs annotated to its term, and untrained, randomly initialized models show most of the agreement before any training. A decoder node can only pass information to the genes of its term, so training on reconstruction can make its activation follow those genes; an untrained model cannot show this, because the fully connected encoder that computes the decoder’s input is untrained as well, so its nodes do not yet carry information about their terms. The same holds for the decoder-based OntoVAE and VEGA, which show no agreement before training and significant agreement after it.

The result should thus not be interpreted as evidence that the activations reveal biology beyond the gene sets from which the network is built. It shows that architecture-derived correspondence is already present before training in the encoder and emerges under the corresponding constraints during training in the decoder, and that GONNECT-SL, whose soft links initially dilute it, recovers much of it in the gene-proximal encoder layers.

More direct validation is limited by the absence of an independent measurement of process activity. One way to address this would be to use a dataset containing perturbation experiments, in which the effect of a gene knockout on a pathway is measured. Alternatively, simulated data could be used, allowing us to define the underlying model of gene expression and process activation and thereby validate GONNECT activations directly.

The signs of the learned weights are unconstrained, so a node’s activation indicates which processes are involved but cannot be read as up- or down-regulation. A sign convention would not remove this limitation: the is_a relations we use state that one term is a subtype of another rather than that one process regulates another, so there is no direction for a weight’s sign to represent. Constraining the weights to be non-negative would instead make each node a strictly additive combination of its inputs, allowing a term’s activation to be attributed to individual genes or child terms. Applied throughout, this would make the network a hierarchical, parts-based decomposition of expression over the ontology, in which a sample’s profile is a sum of process contributions and each process in turn a sum of its constituents, closer in spirit to a biologically structured non-negative matrix factorization than to the autoencoders evaluated here.

The soft-link variant, GONNECT-SL, relaxes the strict GO constraint by allowing connections absent from the ontology to be learned under an L1 penalty. This improves reconstruction beyond the MLP reference while bringing the clustering metrics close to the MLP. The original GO edges retain a weight distribution comparable to the original model, and the soft links remain an order of magnitude smaller, so the prior structure is not simply overwritten. The soft links themselves, however, do not behave as candidate biological relationships. They are unstable across independent training runs, with individual links rarely active in more than one of five runs, and held-out GO edges were not recovered as soft links more often than arbitrary non-GO connections after accounting for local GO degree. Rather than recovering missing biology, the soft links provide the model with additional flexibility to improve reconstruction while largely preserving the original GO structure.

Several limitations stem from the prior itself, and from the processing we apply to it, rather than from the GONNECT architecture. Gene Ontology annotations are unevenly distributed, with well-studied processes, including much of cancer and immune biology, being densely annotated, while less-studied areas remain sparse. The scaffold is therefore richest precisely where the literature is richest. Because the node activations and the enrichment reference are built on the same annotated gene sets, our interpretability comparison also inherits this annotation bias. The structure compounds this. Built from *is_a* relationships, GO is close to a tree in which most terms have a single parent. This favours the encoder over the decoder, for reasons discussed below. More fundamentally, an ontology designed to categorize processes is not necessarily the most appropriate representation for modelling how these processes generate expression patterns. Other priors, such as pathway databases with directed regulatory edges or protein interaction networks, impose different connectivity and would help determine whether the limitations we observe are specific to GO or general to biologically-informed architectures.

A further question is which terms should be included in the first place. Our processing selects terms based on ontology topology alone, while a criterion based on term semantics, for example including or excluding cancer-specific rather than broadly conserved processes, is an orthogonal choice we have not explored. Our processing adds a second source of bias on top of this. Because merging and pruning act where the ontology is sparse, the terms that survive are drawn from its better-annotated regions, and the biology of a merged term is thereafter represented by a broader parent, compounding the annotation bias above.

Selecting the most highly variable genes further conditions the GONNECT architecture on the variation present in the studied cohort. As GO terms are retained based on their connections to the selected input genes, processes represented predominantly by genes with relatively stable expression may be underrepresented or absent from the resulting architecture. We limited this by pruning only redundant paths and merging only where the graph was already too sparse to train on, and saw no sign that the processing drove our results: performance was largely stable across processing settings, except for the GONNECT decoder at the finest settings, and soft-link weights were largely independent of where the ontology was merged or pruned.

Where the prior is placed determines what it costs. A BINN decoder may be too tightly constrained to reconstruct well because each node forces its sibling children to share a single parent activation, leaving them entangled, whereas an encoder node aggregates information from several children rather than reconstructing them. The same constraint makes the decoder configurations markedly slower to optimize, requiring several times as many epochs as the unconstrained reference, whereas the soft-link decoder converges about as fast (Supplementary Table S3). Recovering reconstruction performance therefore requires adding flexibility elsewhere. OntoVAE assigns multiple neurons per GO term, at the cost of several activations per term, which complicates interpretation and requires an aggregation to be chosen before any per-term comparison can be made. GONNECT-SL instead adds soft links, which bring the reconstruction error of the soft-link encoder below that of the unconstrained baseline and its silhouette score to the level of the published BINNs. Taken together, the models we evaluated behave as sparse autoencoders that perform reasonably despite their sparsity, and whose node activations follow their GO terms largely to the extent imposed by the architectural constraints.

Our conclusions are limited to bulk RNA-seq from TCGA tumour samples, but not all findings are expected to be equally dataset-dependent. The encoder/decoder asymmetry is closely related to the near-tree structure of the prior rather than to the data, for the structural reason given above, and we therefore expect its direction to generalize to ontologies with similar characteristics. The soft-link results follow a related argument: if soft links primarily add capacity where the ontology is sparse rather than recover specific missing relationships, their concentration in sparsely connected regions and the lack of a consistently preferred set of links should also be expected in other unevenly annotated priors. By contrast, the limited quantitative benefit of the prior and its near-equivalence to a degree-preserving randomization are more likely to depend on the present dataset, particularly because cancer type dominates expression variance in TCGA.

Normal tissue and single-cell data provide important settings in which to test these findings, with single-cell data being especially informative because process activity may be less diluted by averaging across heterogeneous cell populations. Finally, the architecture itself is not specific to transcriptomics. Any modality that can be linked to an ontology, such as proteomics, could in principle be mapped onto a GONNECT network in the same way. External evaluation across different datasets, biological contexts, and modalities therefore represents an important next step.

## Methods

### Ontology processing

The biological prior knowledge used as model architecture was derived from Gene Ontology (GO)^11^ (monospace>go-basic.obo</monospace>, release 2024-10-27). First, we selected only terms in the *Biological Process* namespace, and only *is_a* relationships, to ensure that the resulting ontology satisfied the properties of a directed acyclic graph (DAG). The ontology graph was extended with the genes from the dataset using the gene annotation file (GAF) for *Homo sapiens* (2024-07-27 release). GO terms without any genes linked to their subtree were removed from the DAG (Figure 6a).

**Figure 6.**
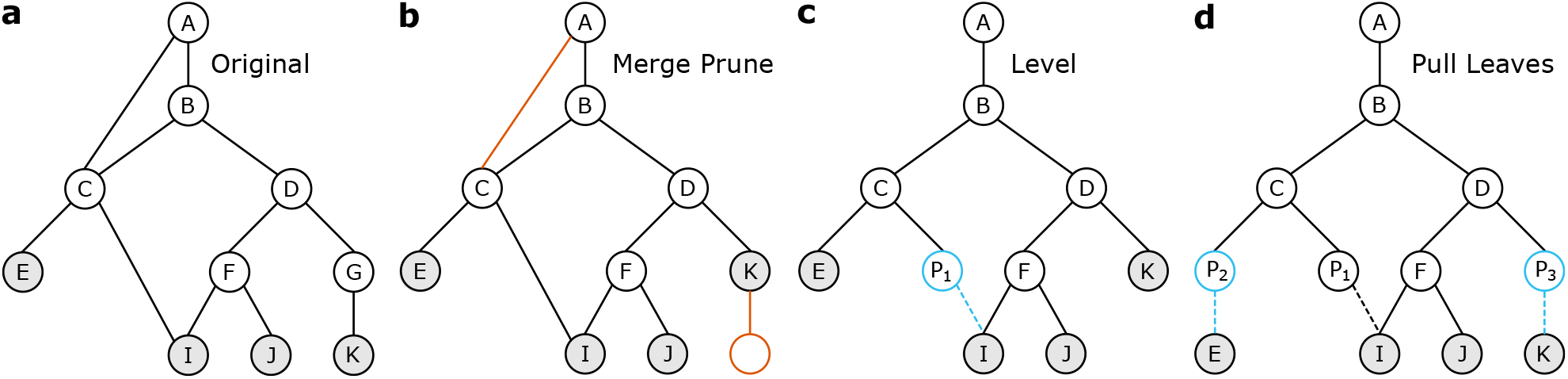
Illustration of the different GO processing steps. Nodes and edges that are coloured red in a panel are removed from the graph during that processing step, those coloured blue are added to the graph. Dotted lines represent edges towards proxy terms. **a**) A graph with similar properties as GO. Nodes *E, I, J* and *K* (coloured) represent genes that have been linked to their annotated GO terms. **b**) Effect of the merge-prune operation for a parent threshold of 1 and a child threshold of 2. Node *G* gets merged into node *D* and the skip connection *AC* gets removed. **c**) The levelling operation ensures that all paths from node *I* to the root have the same length, by introducing a proxy term *P*_1_. **d**) Gene nodes *E* and *K* are moved down by introducing proxy terms *P*_2_ and *P*_3_, such that all genes are at the same depth. Together with the levelling operation, this ensures the resulting graph is suitable for use as a neural network structure.

#### Tree pruning

Terms with low connectivity were removed to reduce the size of the ontology while maintaining its biological interpretability. Based on separate parent and child-term threshold hyperparameters (*pt* and *ct*, set to 1 and 30 respectively), any term with too few parents or children in the hierarchy was merged into its parent(s). This merge operation links all children of the merged term to all parents of the merged term and subsequently removes the term itself from the ontology (Figure 6b). To reduce redundancy, we also prune what we call “skip connections”, where an edge creates a bypass of a term by connecting a node from its subtree to a node in its supertree. We alternatingly apply these merge and prune operations on the tree until convergence, resulting in a more compact, yet still interpretable hierarchy. After these merge and prune iterations, we remove any layer that does not exceed a minimum width (the number of terms in that layer; we used *min* = 50), by merging all terms at that depth into their parents.

#### Neural network compatibility

Each term in the processed ontology is represented by a node in the neural network, and each relationship between two terms is modelled with a learnable weight between the two corresponding nodes. But for a graph to be used as the architecture of an MLP, we require it to be acyclic and balanced. Acyclicity is already ensured by using only *is_a* relationships of GO, which is naturally hierarchical and therefore acyclic. The balanced property refers to the fact that all leaf nodes should be at equal depth, and that all paths from the root to a node have the same length. This is not naturally the case, because even after the initial pruning there is redundancy in the graph, causing what we refer to as “ambiguous node depths” (see node *I* in Figure 6). To ensure the unambiguity of the path lengths, the graph was traversed depth-first and proxy terms were added whenever different paths to a leaf node had different length (Figure 6c). This process was repeated for multiple traversals until the graph converged. To ensure all leaf nodes (corresponding to the genes) were located in the bottom layer, we “pulled” the leaves down by repeatedly inserting proxy terms above genes until each gene reached the maximum depth of the graph (Figure 6d).

The resulting graph has a layout that can be used directly as an MLP without requiring residual connections.

### Gene expression processing

The dataset used for training and evaluation was constructed and downloaded from the Genomic Data Commons (GDC) portal^18^. Samples were included if they were open access, part of the TCGA program, and had available gene expression data.

The retrieved dataset contained 10,498 samples from 9,648 patients, with transcripts per million (TPM) data on 59,427 genes. The TPM values *v*_*i*_ were log-transformed as follows: *v*_*i*_ → log(*v*_*i*_ + 1).

For each gene, the Uniprot ID was retrieved, to link gene expression to an ontology term ^19^. Genes without a match were dropped, after which the gene names of the remaining genes were replaced with their Uniprot ID. The resulting IDs were intersected with the set of Uniprot IDs that have at least one GO-annotation in the Biological Process namespace according to the gene annotation file (GAF). Samples from healthy tissue and genes with zero variance were dropped, leaving 9,797 samples containing 17,491 genes. In light of computational costs, data availability, and model complexity, we trained our models on the 1,000 most highly variable genes in the dataset, selected using the standard Seurat procedure^20^. Each of these genes was then standardized to zero mean and unit variance across samples, so that a mean squared error of 1 corresponds to predicting each gene at its mean.

### GONNECT

The neural network structure derived from GO is implemented by turning the connections between layers of the network into weight matrices. A nonzero element represents an edge between nodes, and thus a relationship between their associated GO terms. Rather than constructing the full weight matrix, we construct two sparse masks per layer (*i*): one for all learnable weights, denoted by **M**_**e**_ and corresponding to biological connections, and one for the proxy terms, denoted by **M**_**p**_. The latter is used to fix the weights leading to proxy terms in the next layer at 1, such that proxy terms introduce no additional learnable weights. This ensures uninterrupted signal propagation through proxy terms and preserves interpretability. The resulting forward pass is defined as:

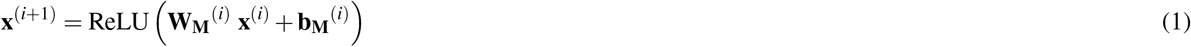

where *i* denotes the network layer, **x** the input and **W**_**M**_ and **b**_**M**_ are the masked versions of the full weight matrix **W** and bias vector **b**, for which the following holds

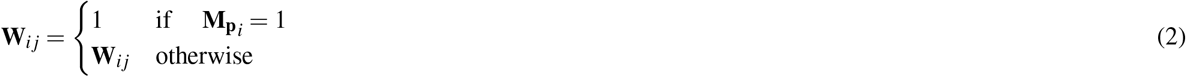

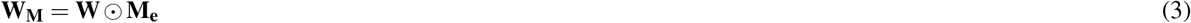

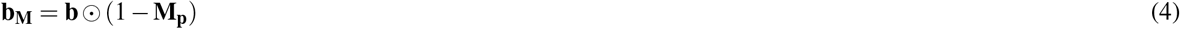

where ⊙ denotes the Hadamard product. At model initialization, the weight matrices of GONNECT are filled by drawing from a Kaiming uniform distribution^21^. After initialization and after each optimization step during training, weight masking is repeated to ensure that the masked values remain fixed.

#### GONNECT-SL

Like original GONNECT, GONNECT-SL stores similar edge masks and proxy masks. After model initialization using a Kaiming uniform distribution, these masks are used to fix the weights towards proxy terms to 1 and proxy biases to 0 (Eqs. 2-4). However, instead of setting non-GO weights to 0, they are initialized as soft links from a normal distribution with mean µ = 0 and standard deviation σ = 0.001. Weights and biases of proxy terms are kept constant by reapplying masks after each optimization step during training.

An additional loss term was included to penalize the absolute value of all soft links, i.e. weights representing edges not present in the ontology graph. The resulting loss function *L*_*SL*_ is used to train these models.

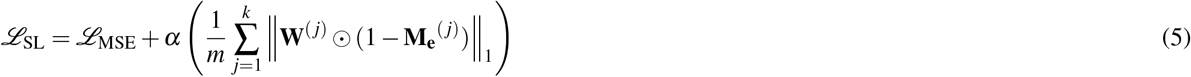

Where α is the regularization factor, **W**^(*j*)^ the weight matrix corresponding to the *j*-th out of *k* network layers, **M**_**e**_^(*j*)^ the edge mask for that layer, ⊙ denotes the Hadamard product, and *m* the total number of soft links. The motivation for weight regularization using the ℓ_1_-norm is to allow some soft links to receive a relatively high weight, while keeping the majority of soft-link weights close to zero. In extremes, a soft-link model where α = 0 is equivalent to a fully connected MLP, while α = ∞ is equivalent to the original ontology-derived GONNECT model. The α hyperparameter was optimized to maximize reconstruction performance while minimizing the number of non-zero soft-link weights. We swept α from 10^2^ to 10^5^ (Supplementary Figures S7 and S8), training each configuration in the sweep for a fixed 1,000 epochs so that the settings are directly comparable. Throughout, a soft link is called *active* when its weight satisfies |*w*| *>* 0.1. The models reported in the main text use α = 1 · 10^2^ and were trained with early stopping.

### Baselines

We also compared GONNECT with two published biologically-informed autoencoders, OntoVAE^14^ and VEGA^15^, using the same repeated random splits as GONNECT. We used each model’s published architecture, optimization, hyperparameters and gene-set definitions without modification; only the input expression data and the train/validation/test splits were replaced by ours.

OntoVAE is a variational autoencoder that wires the GO hierarchy into its decoder, so that each latent dimension and decoder node corresponds to a GO term and the latent space is decoded back to genes through the ontology. To relax the constraint of a single scalar per term, it assigns three neurons to each GO term. Its gene sets were derived from GO using the same processing pipeline as the original OntoVAE implementation.

VEGA is an interpretable variational autoencoder whose masked linear decoder ties each latent dimension to a predefined gene set, using a flat, non-hierarchical prior rather than an ontology. We evaluated it in two configurations, using the gene-set collections from its original publication: the 50 Hallmark gene sets^22^ and the 674 Reactome pathways^23^, both drawn from MSigDB.

### Randomized models

We also compared each ontology-based model against two randomized versions of the hierarchical structure. Randomization was applied independently within each layer. We refer to the degree-preserving variant as GONNECT-DPR and the fully randomized variant as GONNECT-FR throughout, with the analogous variants computed for OntoVAE and VEGA.

GONNECT-DPR preserves every node’s in- and out-degree while discarding edge identity. Within each layer we repeatedly select two edges (*u* → *x*) and (*v* → *y*) and replace them with (*u* → *y*) and (*v* → *x*), rejecting the swap if *u* = *v, x* = *y*, or either new edge already exists. This leaves deg_out_(*u*), deg_out_(*v*), deg_in_(*x*) and deg_in_(*y*) unchanged. We perform *Q* · *E* accepted swaps per layer, where *E* is the number of edges in that layer and *Q* = 100 is a standard mixing heuristic^24^, following XSwap^25^.

GONNECT-FR preserves only the number of edges in each layer. For every layer we count its *E* edges and then draw *E* distinct source–target pairs uniformly at random from all pairs between that layer and the next, discarding the original wiring entirely. Both in- and out-degrees are therefore free to vary and no edge identity is retained, so the layer keeps its sparsity while losing the degree structure that GONNECT-DPR preserves.

### Optimization and evaluation

#### Model optimization

The autoencoder models were trained using the mean squared error (MSE) between input- and reconstructed gene expression data, optimized using stochastic gradient descent with a learning rate of 0.01, a momentum of 0.9, and a batch size of 100. Each model was trained in five independent runs, each with its own random 70%-15%-15% train-validation-test split and weight initialization. The same five splits were reused across models so that comparisons were paired, and all reported metrics were computed on the held-out test split of the corresponding run. Training used early stopping with patience parameter *p* = 10, retaining the model from the final epoch rather than the best-scoring one. The α sweep for GONNECT-SL (Supplementary Figures S7 and S8) was instead run for a fixed 1,000 epochs with early stopping disabled.

#### Metrics

We evaluate each model on four aggregate metrics, computed across five independent training runs. All performance metrics are evaluated on held-out test samples. Reconstruction quality is measured by the mean squared error (MSE) between the input and reconstructed gene expression (lower is better). The other three metrics assess how well the latent space separates the cancer types, using the ground-truth cancer-type labels to define the clusters. We use three because they are sensitive to different things: the silhouette score is geometric and reflects how compact and well separated the clusters are, whereas ARI and NMI depend only on the discrete clustering. A model can therefore score well on one and poorly on the others, as the GONNECT decoder did at low child-term thresholds. Cancer type serves as a proxy for biological structure the models were never trained on, so agreement with it indicates a biologically meaningful latent organization. The silhouette score (SS) compares intra- to inter-cluster distances to capture how compact and well separated the clusters are^26^, ranging from −1 to 1: SS ≈ −1 indicates compact, well separated clusters, SS ≈ 0 for overlapping clusters, and SS ≈ 1 indicates incorrect assignments. The adjusted Rand index (ARI) measures the pairwise agreement between the true labels and an unsupervised *k*-means clustering of the embedding, with *k* set to the number of cancer types^27^. It equals 1 for identical clusterings and is close to 0 for random agreement. Normalized mutual information (NMI) measures the shared information between the same two clusterings^28^, ranging from 0 for no shared information to 1 for perfect agreement. For all three metrics, higher values indicate better performance.

#### k-nearest-neighbour purity

Purity is evaluated on held-out test samples only. For each test sample we retrieve its *k* = 10 nearest neighbours among the other test samples in the latent space by Euclidean distance, and record the fraction of them carrying the same cancer-type label. Purity is the mean of this fraction over all test samples of a cancer type, averaged over the five model instances, ranging from 0 to 1 with higher values indicating less mixing. A cancer type with fewer than *k* test samples cannot reach full purity, so types with fewer than *k* test samples in any of the five splits (some have as few as three test samples) are not shown. Because the cancer types differ strongly in abundance, we additionally report the purity expected when the *k* neighbours are drawn at random from the test samples, which equals the frequency of that cancer type among them and serves as a class-imbalance reference. Because purity depends only on which samples are nearest and not on how far away they are, it measures local label mixing per cancer type rather than the geometry of the embedding as a whole.

#### Latent-space visualization

Latent spaces of all 9,797 samples were visualized in two dimensions with t-SNE^29^, as implemented in scikit-learn, using a perplexity of 30, PCA initialization, at most 1,000 iterations and a fixed random seed; all remaining parameters were left at their defaults. These visualizations are qualitative, and no metric is computed from them.

#### Statistical testing

Models were compared with a paired two-sided *t*-test, with runs paired across the five splits. The reference was the MLP model in Figure 2, the original graph in Figure 3 and Supplementary Figure S4, and the configuration used in the main text in Supplementary Figure S1. The Benjamini–Hochberg procedure was applied for multiple-testing correction within each metric. Multiple significance thresholds are reported: ∗ (*p* ≤ 0.05), ∗∗ (*p* ≤ 0.01) and ∗∗∗ (*p* ≤ 0.001).

### Comparison of node activations with gene-set enrichment

To test whether GONNECT’s GO-term nodes carry the biology their label implies, we related each node’s activation-based cancer-type specificity to a gene-set-level enrichment reference.

First, we identified differentially expressed genes for each cancer type. Every gene was tested with a Wilcoxon rank-sum test comparing its z-scored expression in that cancer type against all samples from all other cancer types pooled. The effect size is the difference of means in z-units. Gene-level *p*-values were Benjamini–Hochberg FDR-adjusted at a 5% FDR.

We then constructed a gene set per GO term, to be able to perform an enrichment analysis. A GO-term node’s gene set is its receptive field: every input gene with a path to the node in the processed GO graph, that is, the genes annotated to that term or to any of its descendants. Because the decoder mirrors the encoder’s wiring, a term’s decoder node projects to the same genes and was assigned the same set. Gene sets were size-filtered to between 3 and 500 genes, which retained 345 of the 623 GO terms, including 86 of the 109 bottleneck terms. Genes were ranked per cancer type by the product of the differential-expression effect size and − log_10_ *p*_adj_, and enrichment was computed with monospace>gseapy.prerank </monospace>(1,000 permutations, weight 1.0, descending rank, sizes 3 to 500, seed 42). The enrichment score carried downstream is −log_10_ of the nominal *p*, floored at 10^−3^ to reflect the resolution of 1,000 permutations. Baseline models were scored analogously. The resulting enrichment scores are then used as the reference against which to compare our own activations.

Interpretability of GONNECT activations is evaluated by their ability to distinguish cancer types. For each node coupled to a GO term we measured how well its single activation distinguishes one cancer type from the rest in a one-versus-rest setup. Per seed, each node’s activation was scored by the area under the ROC curve (ROC-AUC, scikit-learn) using a single-feature logistic regression, and this specificity was carried forward separately for each seed. For the GONNECT models, including the untrained control, specificity was computed on the primary-tumour samples (9,248 samples from 31 cancer types; LAML has no primary-tumour samples). For OntoVAE and VEGA, whose activations are available only for the held-out test split, it was computed on the test-split samples of all sample types.

We then compare this cancer type specificity against the GSEA-derived reference. For each seed and cancer type we computed the Spearman rank correlation between activation-based specificity and GSEA enrichment (− log_10_ nominal *p*) across the GO terms of a layer, requiring at least five terms with both values available, took the median of these correlations over cancer types, and report the mean of this per-seed statistic over the five seeds. To evaluate significance, we use permutation testing in which, for each seed independently, the GO-term columns of the activation matrix are shuffled while the rows are held fixed, breaking the correspondence between activation and enrichment, after which the mean over seeds is recomputed. Across 1,000 permutations, the observed mean is compared to its permuted values through a one-sided permutation *p*-value, *p* = (1 + # *{*permuted ≥ observed})*/*(1 + *n*_perm_). Because the permutation jointly accounts for all terms and cancer types, it serves as the multiple-testing control and no further correction is applied.

The permutation test shuffles term identity, but it does not quantify how much agreement follows from the ontology-derived wiring itself, since every encoder node aggregates the genes annotated to its term irrespective of training. We therefore added an untrained control: the same analysis was repeated on randomly initialized models that were never optimized, using identical layers, gene sets, activation scoring and permutation scheme. The untrained GONNECT, GONNECT-SL and GONNECT-DPR models were built exactly as for training, with the default PyTorch initialization of the linear layers followed by the GO mask, or its degree-preserving randomization for GONNECT-DPR (non-edges set to zero, proxy weights to one) and, for GONNECT-SL, soft links drawn from a normal distribution with mean 0 and standard deviation 10^−3^; the fully connected half of the encoder and decoder models was left untrained as well. We used 50 initializations per model (seeds 0–49), scored on the same primary-tumour samples as the trained models. The untrained OntoVAE and VEGA models were built with each package’s own constructor and default initialization, in the same configuration and on the same ontologies as the trained runs. We used 50 initializations per model and gene-set database, 10 on each of the five data splits, each scored on the test samples of its split like the trained models, with activations taken at the posterior mean. Each initialization was scored separately, and the resulting agreement is reported alongside the trained models in Figure 4.

### Soft-link analyses

#### Seed stability

Here the five seeds are the five runs of the main experiment, which differ in both weight initialization and data split. Across the five seeds, weight reproducibility was quantified per configuration by the pairwise Spearman rank correlation of the full per-edge |*w*| vectors, giving a symmetric correlation matrix. It is accompanied by active-link consistency, the number of seeds (out of five) in which each edge is active at |*w* |*>* 0.1, stacked by layer. Encoder and decoder were summarized separately, with per-layer weight-magnitude densities also reported.

#### Held-out GO-edge recovery

To test whether held-out GO edges are recovered, we trained models in which a random 10% of the ontology edges were held out, and asked whether these removed edges re-acquire higher soft-link weights than edges never present in GO. Analysis of the learned weights was performed in a degree-controlled fashion, to separate any recovery effect from local connectivity.

Each source node’s hard-link degree was binned into log-spaced bins ([1, 2, 4, 8, 16, 32, 64, 128, 256, 512]), and removed and non-removed edges were only ever compared within the same layer and degree bin. Within each bin, the weight magnitudes of removed and non-removed edges were compared with a two-sided Mann–Whitney *U* test, averaged over the five seeds. As an additional check that any signal was not an artefact of bin density, each removed edge was also ranked among the other soft links of its source node.

#### Merge and prune correlation

To test whether soft links concentrate where the ontology was most heavily processed, we related soft-link magnitude to the number of GO-graph edits incident on an edge’s endpoints. Eight edit types were tracked, the combinations of source or sink, sub-tree or super-tree, and prune or merge, across the eight network layers, giving an 8 *×* 8 panel grid. Source counts were restricted to edges with a GO-term source and sink counts to edges with a GO-term sink, since proxy and gene endpoints carry zero edits by construction. Weight was analysed as log_10_ |*w*| against raw edit counts. Each panel statistic is a Spearman rank correlation, subsampled to at most 50,000 edges for speed, with *p*-values Bonferroni-corrected for the 64 tests.

### Implementation details

All models are implemented in Python using pytorch (v 2.5.1). Parsing of the OBO file containing the ontology was performed using the goatools package (v 1.4.12)^30^.

## Acknowledgements

We thank the anonymous reviewers for their constructive comments, which substantially improved the manuscript. This project was funded by BRAINSCAPES, an NWO Gravitation project (grant number 024.004.012).

## Author contributions statement

T.V. and M.L. contributed equally to this work. T.V., M.L. and M.R. conceived the study and designed the experiments. M.L., T.V. and W.M. performed the experiments and analysed the results. T.V. and M.L. wrote the manuscript. M.R. supervised the project and acquired funding. All authors reviewed and approved the final manuscript.

## Code availability

All code used in data processing, model construction and training, and analysis is publicly available at https://github.com/DelftBioinformaticsLab/GONNECT.

## Data availability

All processed data used in this study originates from the TCGA dataset and the Gene Ontology, and is available for download at http://doi.org/10.4121/0d78788b-6bd7-4941-a942-245309107b6d.

## References

1. Sarker, I. H. Machine learning: Algorithms, real-world applications and research directions. SN Comput. Sci. 2, 160, DOI: 10.1007/s42979-021-00592-x (2021).

2. Das, A. & Rad, P. Opportunities and challenges in explainable artificial intelligence (XAI): A survey. arXiv preprint arXiv:2006.11371 DOI: 10.48550/arXiv.2006.11371 (2020).

3. Bell, A., Solano-Kamaiko, I., Nov, O. & Stoyanovich, J. It’s just not that simple: An empirical study of the accuracy-explainability trade-off in machine learning for public policy. In Proceedings of the 2022 ACM Conference on Fairness, Accountability, and Transparency, FAccT ‘22, 248–266, DOI: 10.1145/3531146.3533090 (Association for Computing Machinery, New York, NY, USA, 2022).

4. Assis, A., Dantas, J. & Andrade, E. The performance-interpretability trade-off: a comparative study of machine learning models. J. Reliab. Intell. Environ. 11, 1, DOI: 10.1007/s40860-024-00240-0 (2025).

5. Chen, V. et al. Applying interpretable machine learning in computational biology—pitfalls, recommendations and opportunities for new developments. Nat. Methods 21, 1454–1461, DOI: 10.1038/s41592-024-02359-7 (2024).

6. Sidak, D., Schwarzerová, J., Weckwerth, W. & Waldherr, S. Interpretable machine learning methods for predictions in systems biology from omics data. Front. Mol. Biosci. 9, 926623, DOI: 10.3389/fmolb.2022.926623 (2022).

7. Ma, J. et al. Using deep learning to model the hierarchical structure and function of a cell. Nat. Methods 15, 290–298, DOI: 10.1038/nmeth.4627 (2018).

8. Hao, J., Kim, Y., Kim, T. K. & Kang, M. PASNet: Pathway-associated sparse deep neural network for prognosis prediction from high-throughput data. BMC Bioinforma. 19, 510, DOI: 10.1186/s12859-018-2500-z (2018).

9. Elmarakeby, H. A. et al. Biologically informed deep neural network for prostate cancer discovery. Nature 598, 348–352, DOI: 10.1038/s41586-021-03922-4 (2021).

10. Hao, Y., Romano, J. D. & Moore, J. H. Knowledge-guided deep learning models of drug toxicity improve interpretation. Patterns 3,100565, DOI: 10.1016/j.patter.2022.100565 (2022).

11. Ashburner, M. et al. Gene Ontology: tool for the unification of biology. Nat. Genet. 25, 25–29, DOI: 10.1038/75556 (2000).

12. Lotfollahi, M. et al. Biologically informed deep learning to query gene programs in single-cell atlases. Nat. Cell Biol. 25, 337–350, DOI: 10.1038/s41556-022-01072-x (2023).

13. Kuenzi, B. M. et al. Predicting drug response and synergy using a deep learning model of human cancer cells. Cancer Cell 38, 672–684.e6, DOI: 10.1016/j.ccell.2020.09.014 (2020).

14. Doncevic, D. & Herrmann, C. Biologically informed variational autoencoders allow predictive modeling of genetic and drug-induced perturbations. Bioinformatics 39, btad387, DOI: 10.1093/bioinformatics/btad387 (2023).

15. Seninge, L., Anastopoulos, I. N., Ding, H. & Stuart, J. M. VEGA is an interpretable generative model for inferring biological network activity in single-cell transcriptomics. Nat. Commun. 12, 5684, DOI: 10.1038/s41467-021-26017-0 (2021).

16. Caranzano, I. et al. Sparsity is all you need: Rethinking biological pathway-informed approaches in deep learning. arXiv preprint arXiv:2505.04300 DOI: 10.48550/arXiv.2505.04300 (2025).

17. Tomczak, K., Czerwinska, P. & Wiznerowicz, M. The Cancer Genome Atlas (TCGA): an immeasurable source of knowledge. Contemp. Oncol. 19, 68–77, DOI: 10.5114/wo.2014.47136 (2015).

18. Jensen, M. A., Ferretti, V., Grossman, R. L. & Staudt, L. M. The NCI Genomic Data Commons as an engine for precision medicine. Blood 130, 453–459, DOI: 10.1182/blood-2017-03-735654 (2017).

19. The UniProt Consortium. UniProt: the universal protein knowledgebase in 2025. Nucleic Acids Res. 53, D609–D617, DOI: 10.1093/nar/gkae1010 (2024).

20. Satija, R., Farrell, J. A., Gennert, D., Schier, A. F. & Regev, A. Spatial reconstruction of single-cell gene expression data. Nat. Biotechnol. 33, 495–502, DOI: 10.1038/nbt.3192 (2015).

21. He, K., Zhang, X., Ren, S. & Sun, J. Delving deep into rectifiers: Surpassing human-level performance on ImageNet classification. In Proceedings of the 2015 IEEE International Conference on Computer Vision (ICCV), ICCV ‘15, 1026–1034, DOI: 10.1109/ICCV.2015.123 (IEEE Computer Society, USA, 2015).

22. Liberzon, A. et al. The Molecular Signatures Database Hallmark gene set collection. Cell Syst. 1, 417–425, DOI: 10.1016/j.cels.2015.12.004 (2015).

23. Croft, D. et al. Reactome: a database of reactions, pathways and biological processes. Nucleic Acids Res. 39, D691–D697, DOI: 10.1093/nar/gkq1018 (2011).

24. Espinoza, M. On network randomization methods: A negative control study. Tech. Rep., Fairfield University, Fairfield, CT (2012).

25. Hanhijärvi, S., Garriga, G. C. & Puolamäki, K. Randomization techniques for graphs. In Proceedings of the 2009 SIAM International Conference on Data Mining (SDM), 780–791, DOI: 10.1137/1.9781611972795.67 (2009).

26. Rousseeuw, P. J. Silhouettes: a graphical aid to the interpretation and validation of cluster analysis. J. Comput. Appl. Math. 20, 53–65, DOI: 10.1016/0377-0427(87)90125-7 (1987).

27. Hubert, L. & Arabie, P. Comparing partitions. J. Classif. 2, 193–218, DOI: 10.1007/BF01908075 (1985).

28. Lancichinetti, A., Fortunato, S. & Kertész, J. Detecting the overlapping and hierarchical community structure in complex networks. New J. Phys. 11, 033015, DOI: 10.1088/1367-2630/11/3/033015 (2009).

29. van der Maaten, L. & Hinton, G. Visualizing data using t-SNE. J. Mach. Learn. Res. 9, 2579–2605 (2008).

30. Klopfenstein, D. V. et al. GOATOOLS: A Python library for Gene Ontology analyses. Sci. Reports 8, 10872, DOI: 10.1038/s41598-018-28948-z (2018).

