## Supplementary Figures for "A critical evaluation of Gene Ontology priors in biologically-informed neural networks"

---

### Contents

**Supplementary Figure S1.** Extended version of Figure 2 across the GO-processing hyperparameter sweep.

**Supplementary Figure S2.** t-SNE visualizations of all soft-link equivalents of the models shown in Figure 2.

**Supplementary Figure S3.** Per-cancer-type embedding quality for all ten model variants with saved latent spaces.

**Supplementary Figure S4.** Comparison of each model with two randomized alternatives (ARI and NMI).

**Supplementary Figure S5.** Mean activations per cancer type of 20 selected GO terms, GONNECT encoder.

**Supplementary Figure S6.** Mean activations per cancer type of 20 selected GO terms, GONNECT decoder.

**Supplementary Figure S7.** Loss curves of GONNECT-SL training for different values of the soft-link hyperparameter $\alpha$.

**Supplementary Figure S8.** Weight distributions for different values of the soft-link hyperparameter $\alpha$.

**Supplementary Figure S9.** Stability of GONNECT-SL soft-link weights across the five training seeds.

**Supplementary Figure S10.** Held-out GO-edge recovery in the GONNECT-SL encoder, by layer.

**Supplementary Figure S11.** Held-out GO-edge recovery in the GONNECT-SL decoder, by layer.

**Supplementary Figure S12.** Soft-link weight magnitude versus GO-edit count, by edit type and layer.

*Supplementary Tables S1–S5 are provided as separate files.*

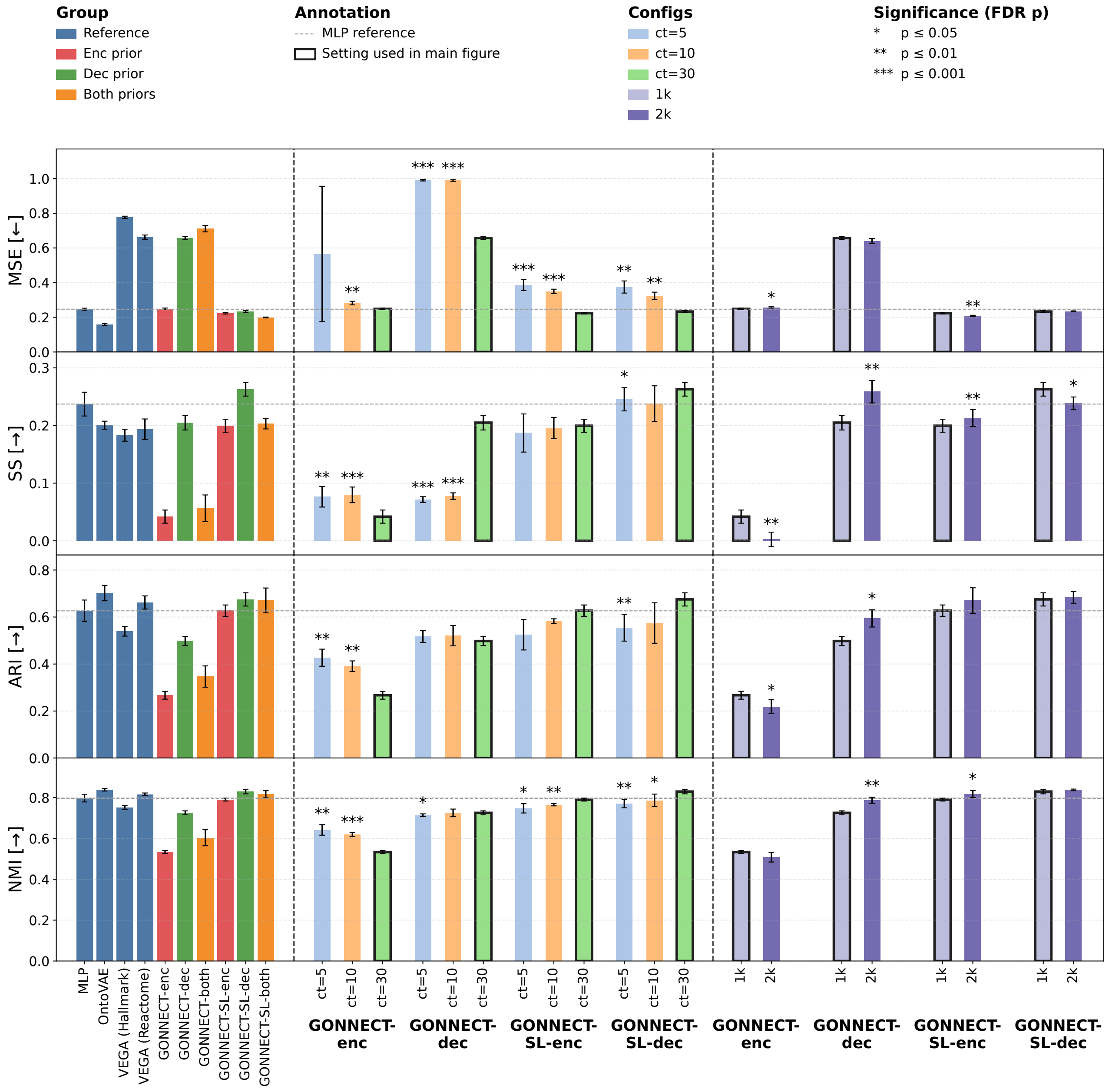


**Supplementary Figure S1.** Extended version of Figure 2 across the GO-processing hyperparameter sweep. (Rows) Mean squared error (MSE), silhouette score (SS), adjusted Rand index (ARI), and normalized mutual information (NMI) for all model variants, computed on the held-out test split. The three column blocks show, from left to right, the models at the settings used in the main text, a child-term threshold sweep (ct=5, 10, 30) at 1,000 input genes, and a comparison of 1,000 against 2,000 input genes at ct=30. The remaining processing parameters were held at their defaults. The configuration used in the main text (ct=30, 1,000 genes) is indicated by a thickened bar outline. Bar height denotes the mean over five independently trained model instances and error bars denote the standard deviation. Significance is reported with respect to the outlined configuration from the main text, and calculated with Benjamini–Hochberg correction (* $p \leq 0.05$, ** $p \leq 0.01$, *** $p \leq 0.001$).

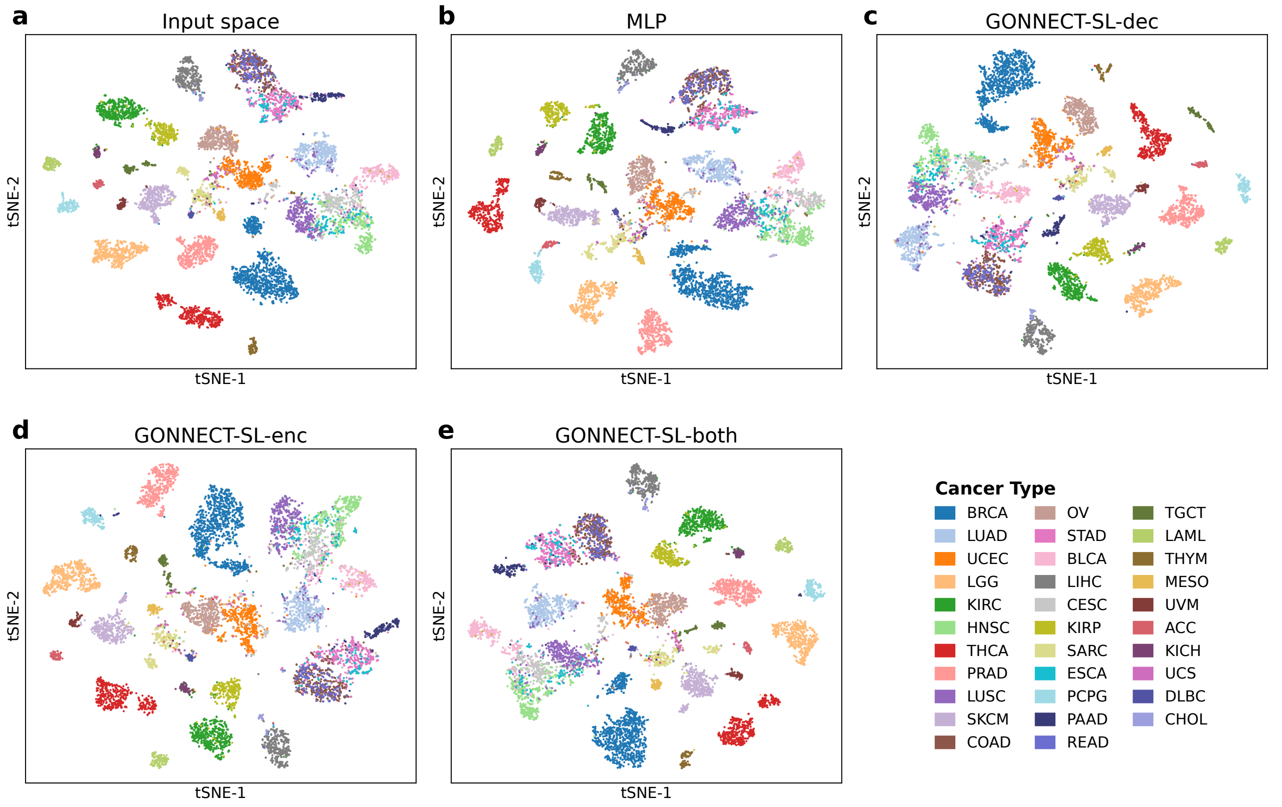


**Supplementary Figure S2.** t-SNE visualizations of all soft-link equivalents of the models shown in Figure 2.

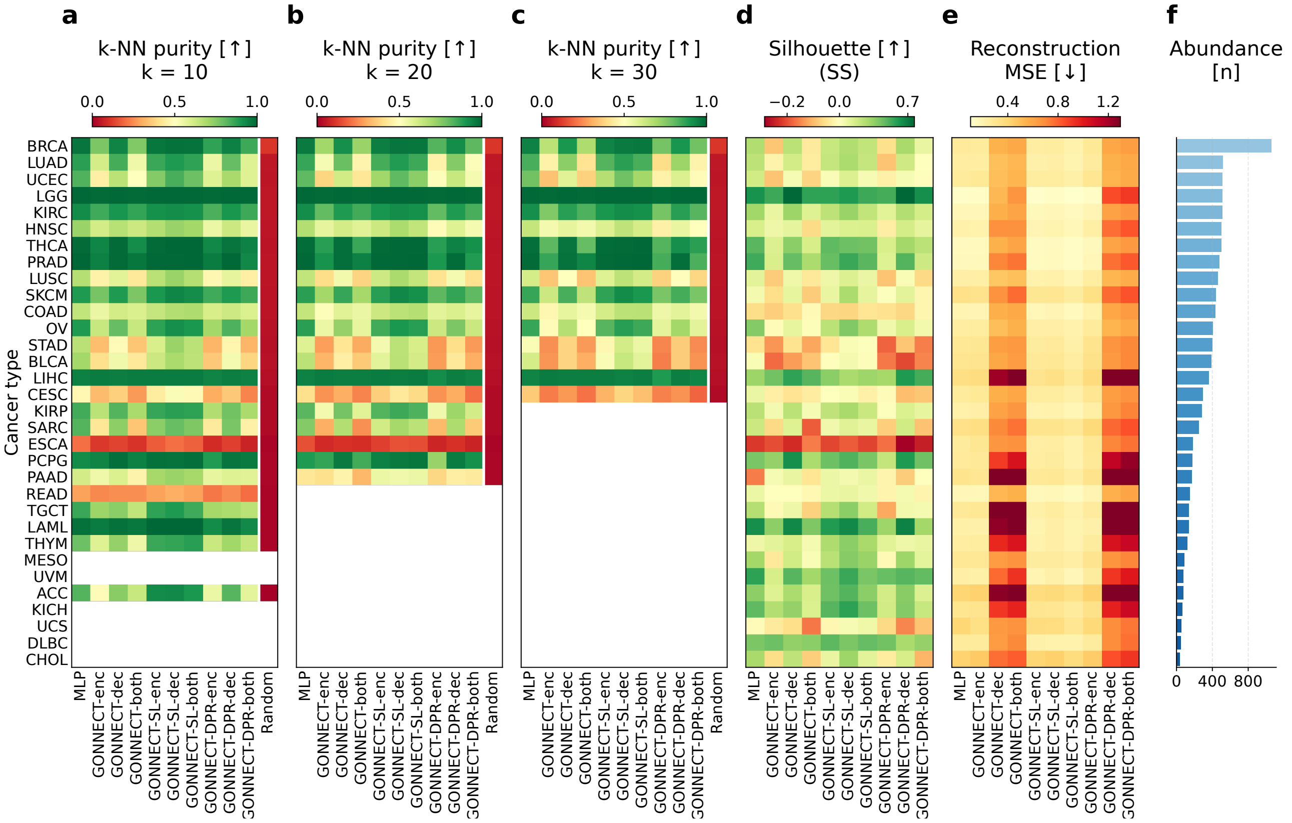


**Supplementary Figure S3.** Per-cancer-type embedding quality for all ten model variants with saved latent spaces, including the degree-preserving random (DPR) controls that the main text reports only in aggregate. Rows are the 32 TCGA cancer types, sorted by abundance (panel **f**); every heatmap carries the same ten models in the same column order. All values are computed on the held-out test split and averaged over five independently trained model instances, and arrows indicate whether the metric improves by ascending or descending in value.

**a–c**) $k$-nearest-neighbour purity, defined in Figure 2 (test samples against their nearest neighbours among the other test samples), at $k = 10$, 20 and 30; comparing the three shows whether a type stays pure as the neighbourhood grows. Cancer types with fewer than $k$ test samples in any of the five splits are not shown. The rightmost column of each purity panel shows the purity expected for randomly drawn neighbours given the class imbalance (Methods). Panel **a** is the metric shown in Figure 2**k**.

**d**) silhouette score, which unlike purity is geometric: it rewards clusters that are both compact and well separated, and turns negative where a type's samples are on average closer to the samples of another type than to those of their own. The colour scale is centred on zero.

**e**) reconstruction MSE, the only panel where lower is better and the only metric not using the cancer type labels.

Purity uses a fixed 0–1 colour scale; silhouette and MSE use data-driven limits.

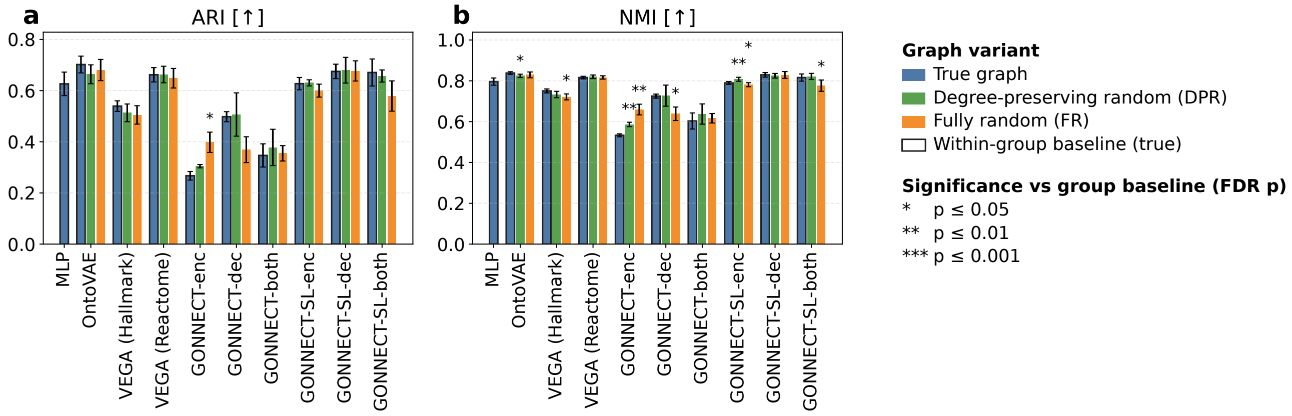


**Supplementary Figure S4.** Comparison of each model with two randomized alternatives, one degree-preserving where each term keeps its degree, but connectivity to other terms is randomized, and one fully randomized where only the number of edges per layer is preserved. Significance is reported with respect to the original model version. We report adjusted Rand index (ARI, panel **a**), and normalized mutual information (NMI, panel **b**), both computed on the held-out test split.

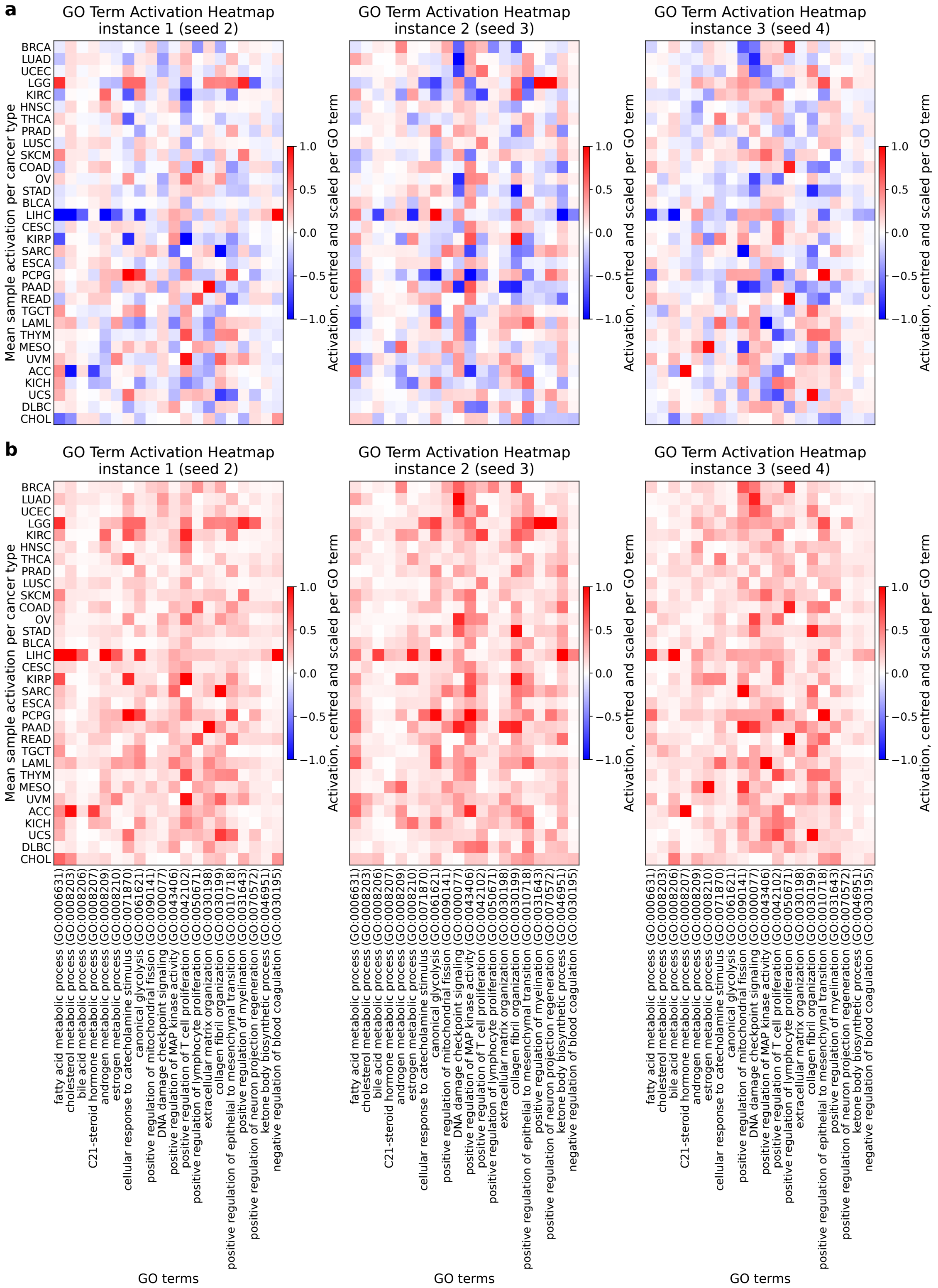


**Supplementary Figure S5.** Mean activations per cancer type of 20 GO terms selected as processes expected to vary in activity across cancer types, computed over all 9,797 samples. The three heatmaps per panel show three instances of a GONNECT encoder model. **a**) Signed activations per term, per cancer type. **b**) Absolute values of the mean activations in panel **a**. Panel **a** shows how the sign of the mean activations appears random across model instances. Panel **b** shows how the magnitudes of some term–cancer type pairs are consistently high across different encoder instances. The complete list of GO terms used in GONNECT-enc, with their per-cancer-type association scores (ROC-AUC), is given in Supplementary Table S4.

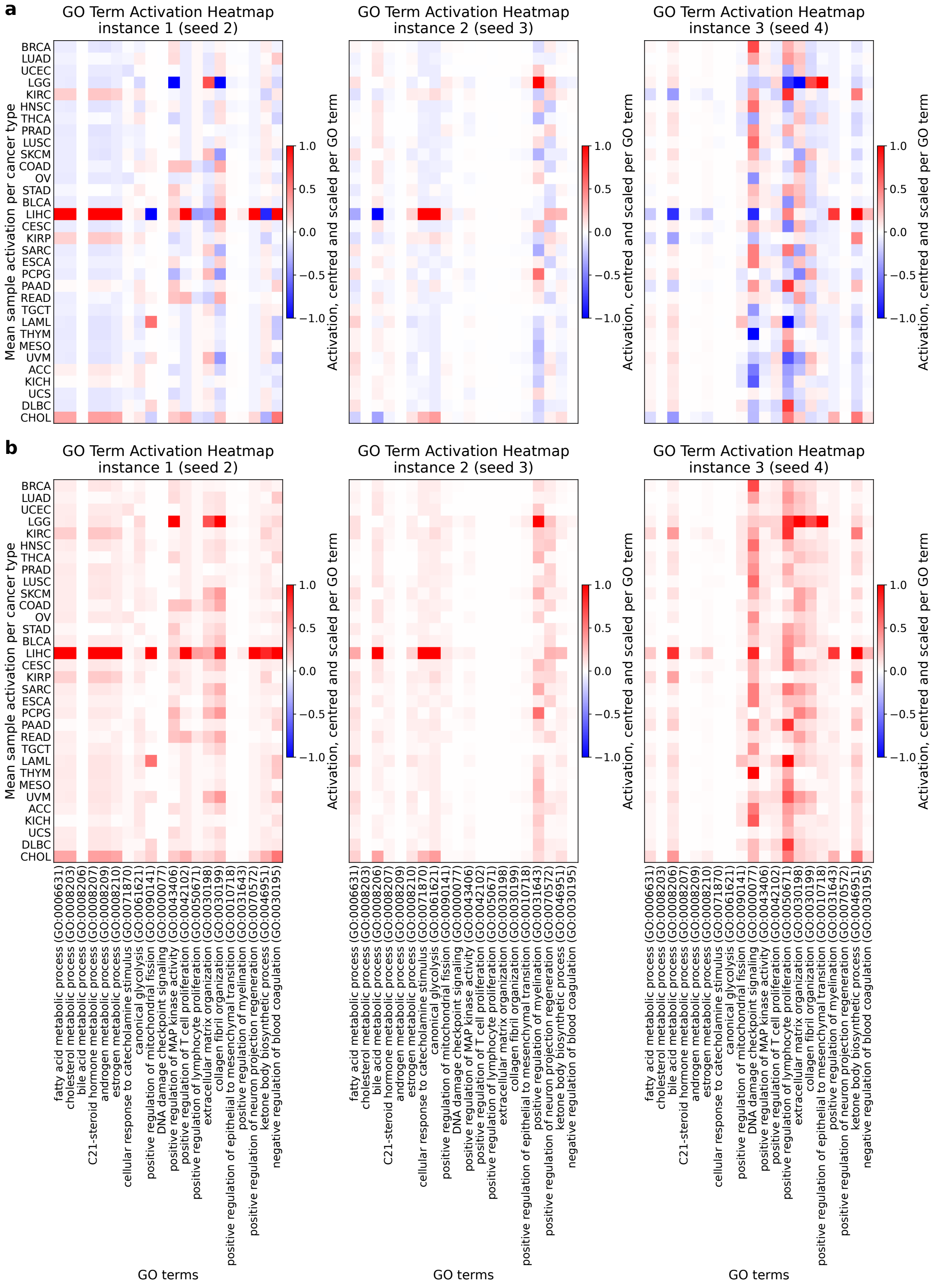


**Supplementary Figure S6.** Mean activations per cancer type of 20 GO terms selected as processes expected to vary in activity across cancer types, computed over all 9,797 samples. The three heatmaps per panel show three instances of a GONNECT decoder model. **a**) Signed activations per term, per cancer type. **b**) Absolute values of the mean activations in panel **a**. Panel **a** shows how the sign of the mean activations appears random across model instances. Panel **b** shows that, unlike in the encoder (Supplementary Figure S5), few term–cancer type pairs are consistently high across decoder instances; the clearest exception is LIHC with liver-related metabolic terms. The complete list of GO terms used in GONNECT-dec, with their per-cancer-type association scores (ROC-AUC), is given in Supplementary Table S5.

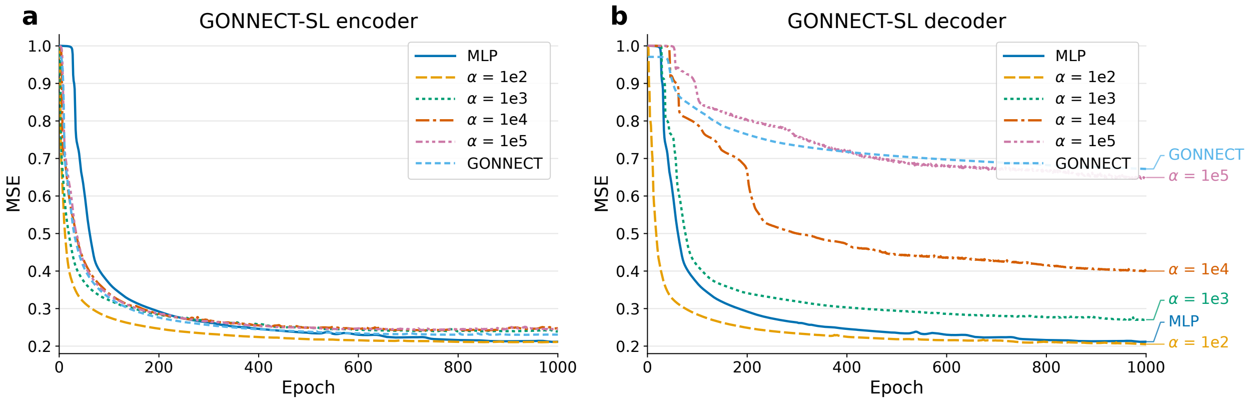


**Supplementary Figure S7.** Loss curves of GONNECT-SL training with different values for soft-link hyperparameter $\alpha$. The models are being trained with a specialized loss function, however, the figure shows regular mean squared error (MSE) of input reconstruction. **a**) Loss curve of a GONNECT-SL encoder module. **b**) Loss curve of a GONNECT-SL decoder module. The larger $\alpha$ imposes stronger regularization on the soft links. This results in fewer active soft links, which is favourable for interpretability, but reduced reconstruction performance. The effect on reconstruction performance is more pronounced for the GONNECT-SL decoder than for the encoder. Each configuration in this sweep was trained for a fixed 1,000 epochs with early stopping disabled, unlike the models reported in the main text.

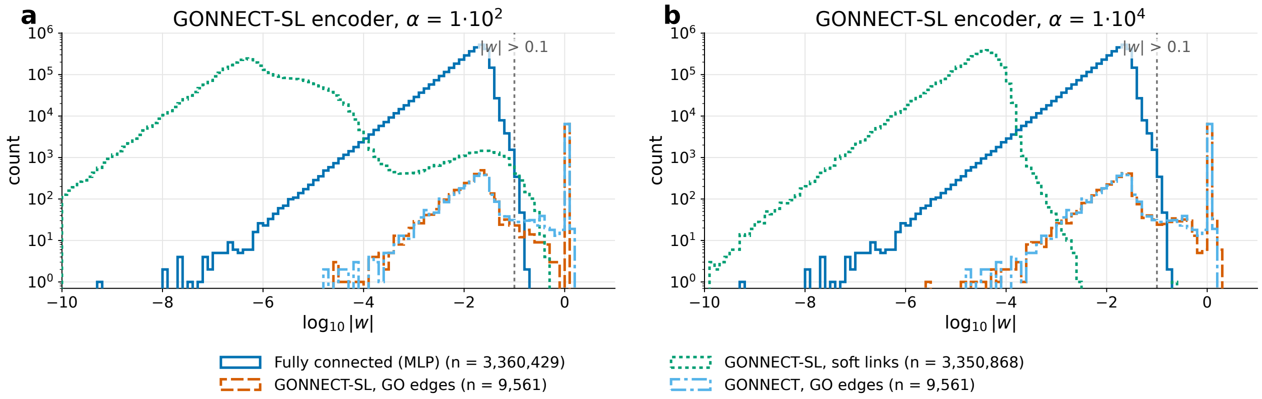


**Supplementary Figure S8.** Weight distributions for different values of soft-link hyperparameter $\alpha$. For comparison, the weight distribution of a fully connected MLP, and the distribution of original GONNECT, are shown. The weights of GONNECT-SL are split up into the weights of links that were already present in GO and those that are not, meaning they can become active soft links. The dotted vertical line marks the $|w| > 0.1$ threshold used to count a soft link as active. As in Supplementary Figure S7, each configuration in this sweep was trained for a fixed 1,000 epochs with early stopping disabled. **a**) Weight distribution for $\alpha = 1 \cdot 10^2$. The relatively low value means that a lot of soft links are active, and most active soft links have relatively small magnitudes. Under this fixed 1,000-epoch schedule the original GO links also lose their high magnitude; the models reported in the main text use the same $\alpha$ but stop early, and retain the GO-weight distribution shown in Figure 5a. **b**) Weight distribution for $\alpha = 1 \cdot 10^4$. The high value means that just one soft link remains active in the encoder, and the model essentially becomes equivalent to the original ontology-based GONNECT model.

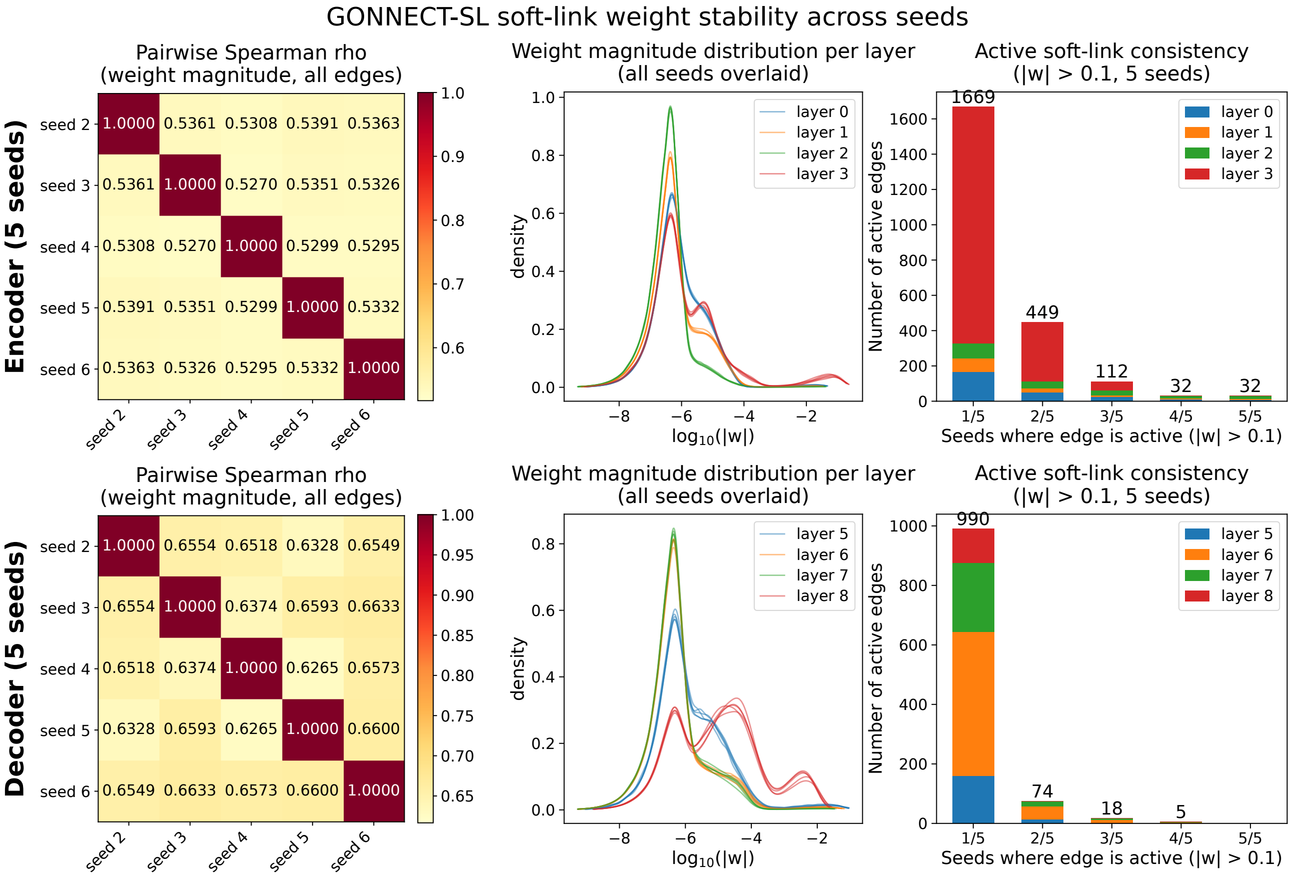


**Supplementary Figure S9.** Stability of GONNECT-SL soft-link weights across the five training seeds, shown separately for the encoder (top) and decoder (bottom). Left: pairwise Spearman rank correlation of the per-edge weight magnitudes $|w|$ between seeds. Middle: per-layer density of weight magnitudes ($\log_{10}|w|$), all seeds overlaid. Right: active soft-link consistency, the number of edges active at $|w| > 0.1$ in a given number of seeds (1 of 5 to 5 of 5), stacked by layer. Most active soft links appear in only one seed, with 32 edges active in all five seeds in the encoder and none in the decoder.

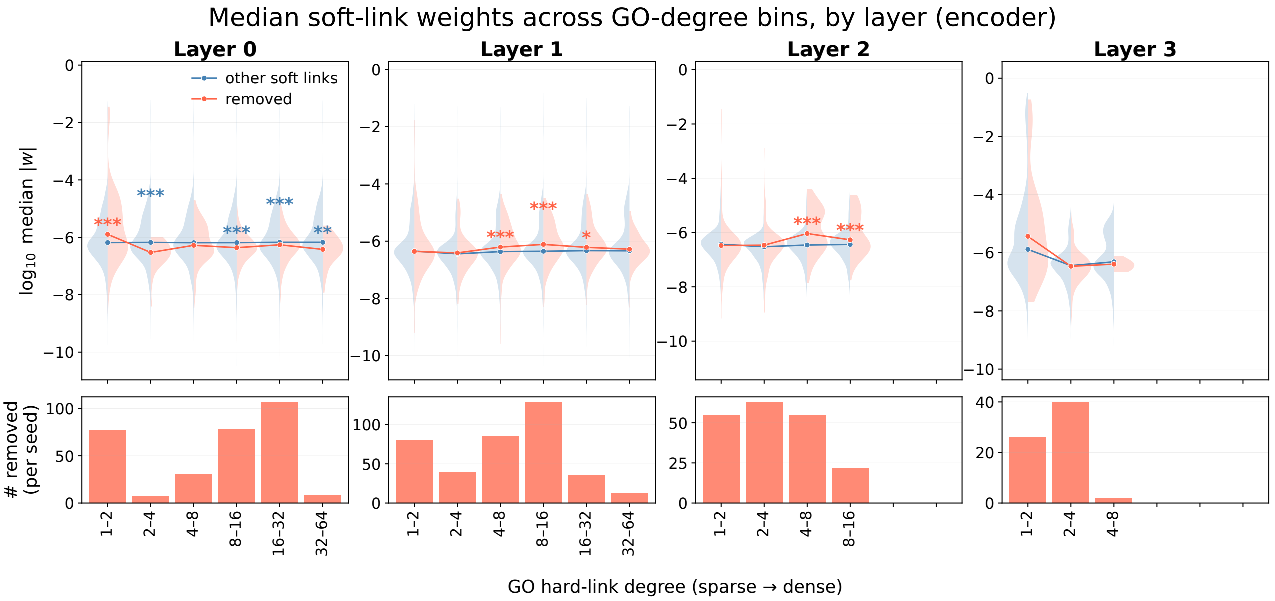


**Supplementary Figure S10.** Held-out GO-edge recovery in the GONNECT-SL encoder, by layer (L0 to L3). For each layer, soft-link weight magnitude ($\log_{10}$ median $|w|$) is compared between edges held out from GO (removed) and other non-GO soft links, binned by the source node's hard-link degree in the ontology (sparse to dense, left to right). Points and lines give the per-bin median over the five seeds and violins show the distribution. Removed and non-removed edges are compared only within the same layer and degree bin (two-sided Mann–Whitney U, averaged over seeds; * $p \leq 0.05$, ** $p \leq 0.01$, *** $p \leq 0.001$). Bottom histograms give the number of removed edges per seed in each bin. Held-out GO edges are not recovered preferentially once local GO degree is accounted for; recovery tracks the density of GO terms in the node's neighbourhood.

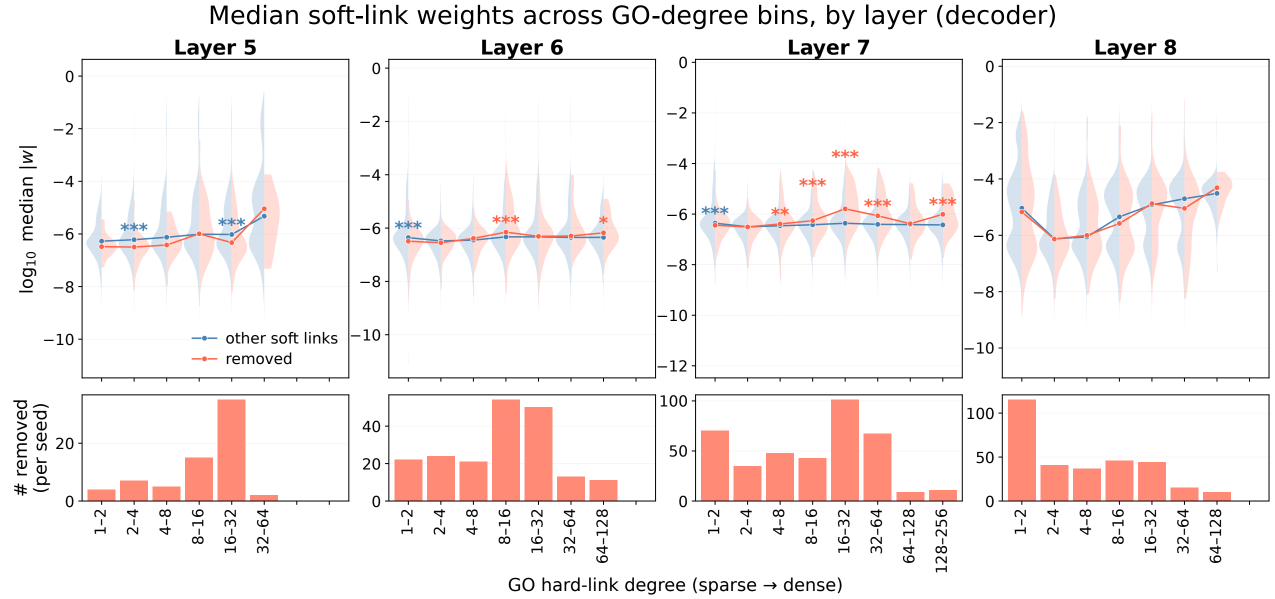


**Supplementary Figure S11.** Held-out GO-edge recovery in the GONNECT-SL decoder, by layer (L5 to L8). As in Supplementary Figure S10, but for the decoder. Soft-link weight magnitude ($\log_{10}$ median $|w|$) is compared between held-out GO edges (removed) and other non-GO soft links within matched layer and source-degree bins (two-sided Mann–Whitney U, averaged over the five seeds; * $p \leq 0.05$, ** $p \leq 0.01$, *** $p \leq 0.001$), with the number of removed edges per seed shown below. As in the encoder, held-out GO edges are not recovered preferentially after controlling for local GO degree.

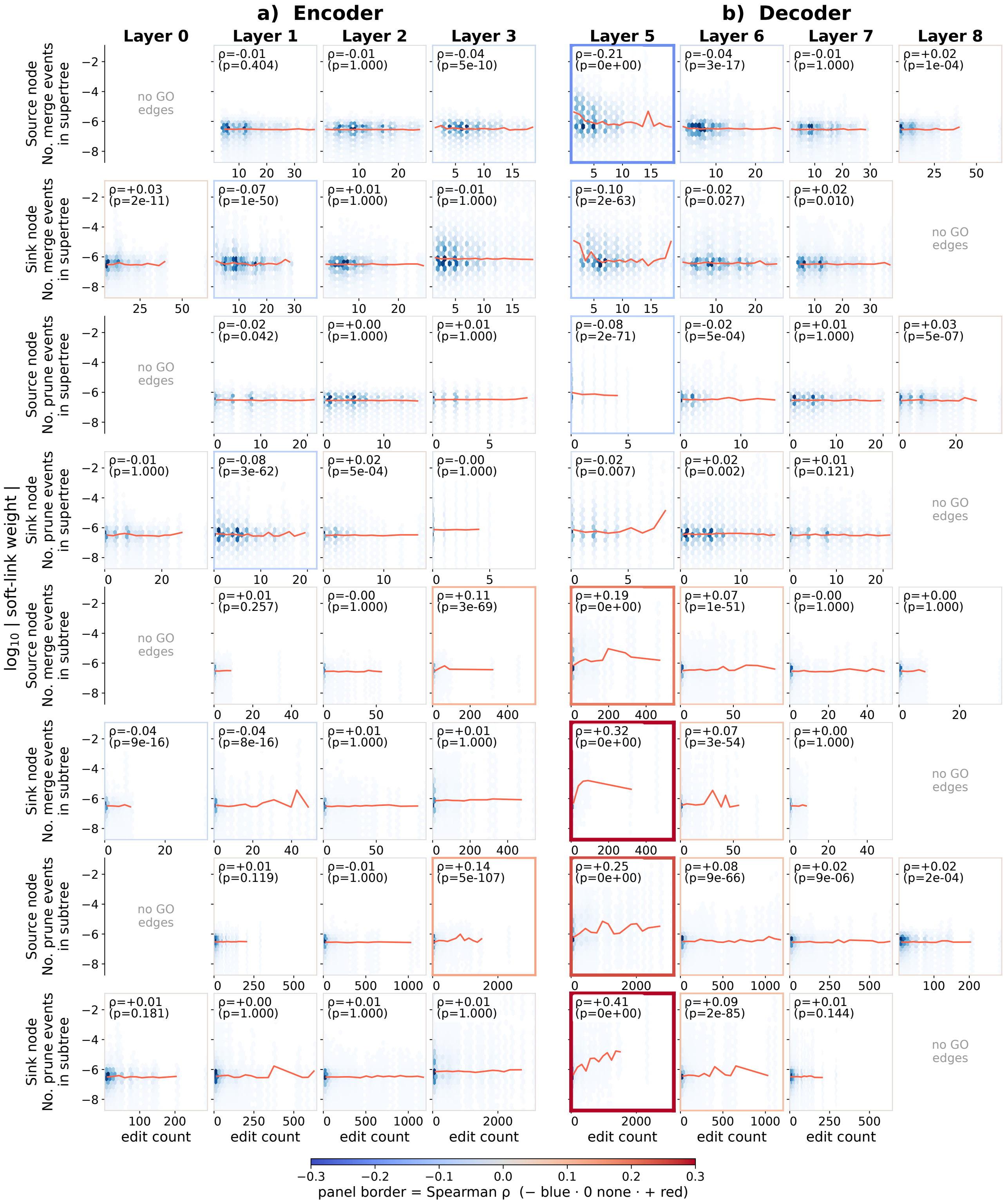


**Supplementary Figure S12. Soft-link weight magnitude versus GO-edit count, by edit type and layer**, for the **(a)** GONNECT-SL-enc and **(b)** GONNECT-SL-dec; columns are layers. Rows are the eight edit types (source/sink × merge/prune × super-/subtree); each panel plots edit count against $\log_{10}|w|$ as a density hexbin with a running median (red line). Only soft links with a GO-term endpoint are shown. Panels are annotated in text, and are also coloured by the Bonferroni-corrected Spearman correlation $\rho$ (blue/white/red).
